# Covalent 14-3-3 Molecular Glues and Heterobifunctional Molecules Against Nuclear Transcription Factors and Regulators

**DOI:** 10.1101/2023.11.06.565850

**Authors:** Qian Shao, Tuong Nghi Duong, Inji Park, Daniel K. Nomura

## Abstract

14-3-3 proteins have the unique ability to bind and sequester a multitude of diverse phosphorylated signaling proteins and transcription factors. Many previous studies have shown that 14-3-3 interactions with specific phosphorylated substrate proteins can be enhanced through small-molecule natural product or fully synthetic molecular glue interactions. However, enhancing 14-3-3 interactions with both therapeutically intractable transcription factor substrates as well as potential neo-substrates to sequester and inhibit their function has remained elusive. One of the 14-3-3 proteins, 14-3-3σ or SFN, has a cysteine C38 at the substrate binding interface near sites where previous 14-3-3 molecular glues have been found to bind. In this study, we screened a fully synthetic cysteine-reactive covalent ligand library to identify molecular glues that enhance interaction of 14-3-3σ with not only druggable transcription factors such as estrogen receptor (ERα), but also challenging oncogenic transcription factors such as YAP and TAZ that are part of the Hippo transducer pathway. We identified a hit EN171 that covalently targets 14-3-3 to enhance 14-3-3 interactions with ERα, YAP, and TAZ leading to impaired estrogen receptor and Hippo pathway transcriptional activity. We further demonstrate that EN171 could not only be used as a molecular glue to enhance native protein interactions, but also could be used as a covalent 14-3-3 recruiter in heterobifunctional molecules to sequester nuclear neo-substrates such as BRD4 into the cytosol. Overall, our study reveals a covalent ligand that acts as a novel 14-3-3 molecular glue for challenging transcription factors such as YAP and TAZ and also demonstrates that these glues can be potentially utilized in heterobifunctional molecules to sequester nuclear neo-substrates out of the nucleus and into the cytosol to enable targeted protein localization.

## Introduction

14-3-3 proteins regulate a vast network of proteins in the cell by mostly recognizing phosphorylated serine or threonine bearing peptides on specific protein substrates. This binding often sequesters these proteins into the cytosol from either a cellular compartments where they may be active or from other protein-protein interactions to modulate their functions ^1–3^. Substrates of 14-3-3 include many important proteins involved in cancer signaling or transcription, including Raf-1, Bad, Bax, Cdc25, Akt, and SOS, as well as transcription factors such as estrogen receptor (ERα), YAP, and TAZ ^1–3^.

Seminal studies by Christian Ottmann, Michelle Arkin, and others have led to the discovery of natural products such as fusicoccin A as well as fully synthetic small-molecules that act as molecule glues by binding to the substrate cleft and enhancing interactions of 14-3-3 with native phosphorylated substrate proteins ^1–12^. While these molecules have enhanced and stabilized interactions between 14-3-3 and several different protein substrates including ERα, ERR*γ*, c-RAF, FOXO1, USP8, and SOS1^1–12^, small-molecule glues have not yet been able to access some of the more therapeutically intractable nuclear transcription factors such as YAP and TAZ in the Hippo transducer pathway. Furthermore, it is not clear whether 14-3-3 molecular glues would be capable of sequestering neo-substrates from their native cellular compartments or protein complexes for potential therapeutic applications.

In this study, we screened a library of cysteine-reactive covalent ligands to exploit a cysteine positioned in the substrate binding cleft of a particular 14-3-3 isoform, 14-3-3σ or SFN, to identify a covalent molecular glue stabilizer of 14-3-3 interactions not only with more druggable targets such as ERα, but also less druggable transcription factors such as YAP and TAZ. We also demonstrated that our covalent 14-3-3 molecular glue stabilizer could be used as a 14-3-3 recruiter in heterobifunctional molecules to sequester nuclear neo-substrate targets out of the nucleus and into the cytosol.

## Results

### Covalent Ligand Screen to identify 14-3-3 Molecular Glue Stabilizers

To identify a covalent 14-3-3 molecular glue stabilizer, we performed a fluorescence polarization screen searching for compounds that would stabilize pure human 14-3-3σ protein and the phosphorylated ERα peptide substrate of 14-3-3 **(Figure 1a; Table S1)**. We identified one major hit compound EN171 that showed robust 14-3-3σ stabilization with phosphorylated ERα peptide to the same extent as the positive control fusicoccin A **(Figure 1a-1b)**. We observed dose-responsive stabilization of 14-3-3σ with ERα with a 50 % effective concentration (EC_50_) of 9 μM. Quite intriguingly, we also observed dose-responsive stabilization of 14-3-3σ with YAP and TAZ phosphopeptides as well with EC_50_ values of 45 and 107 μM, respectively **(Figure 1c)**. To show that this stabilization was depending on 14-3-3, we also showed attenuation of EN171-mediated stabilization between 14-3-3 and ERα, YAP, or TAZ with the 14-3-3 inhibitor R18 **(Figure 1d)**.

**Figure 1.**
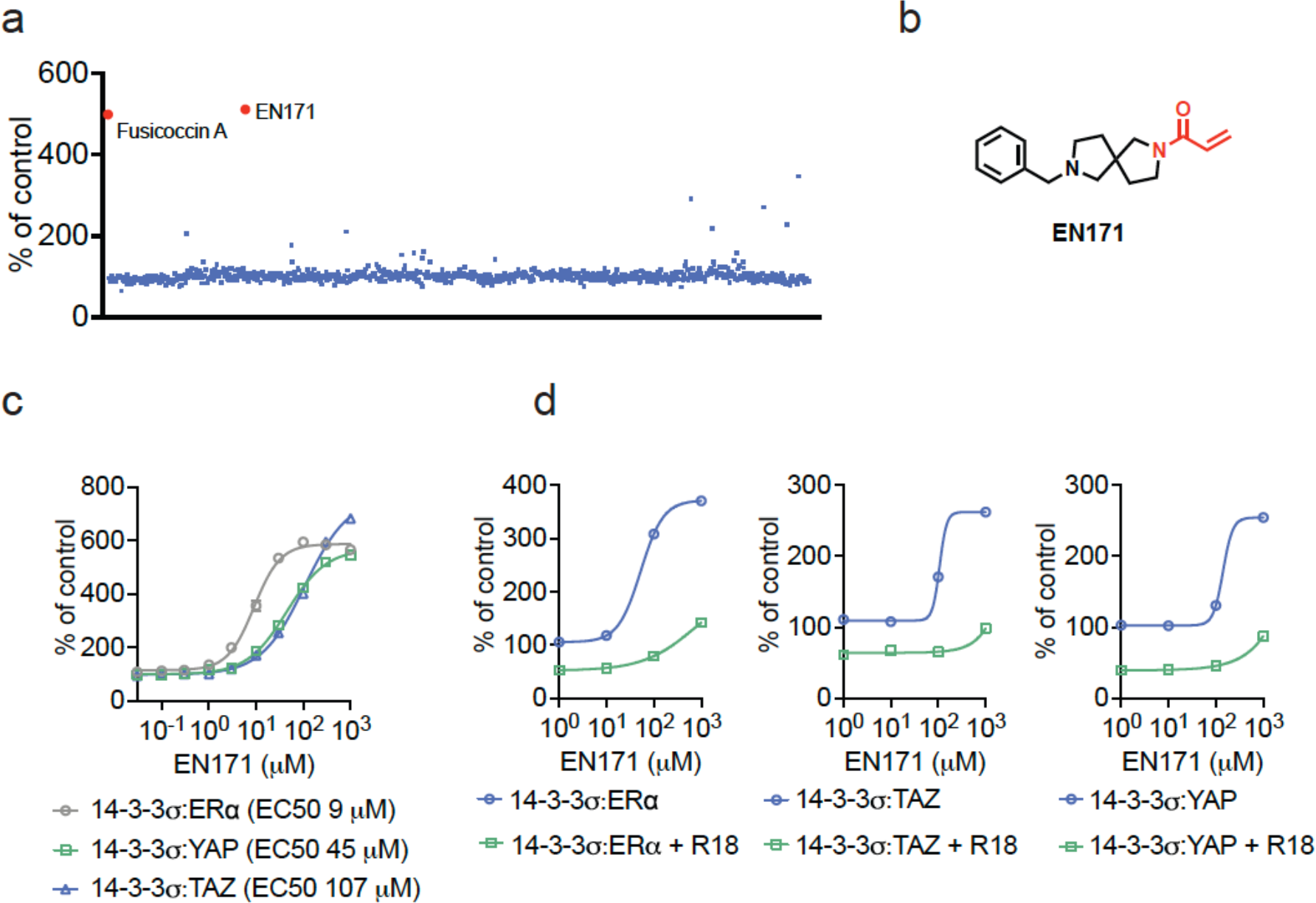
Identification of 14-3-3 molecular glue stabilizers. **(a)** Fluorescence polarization (FP) screening of a cysteine-reactive covalent ligand library (125 μM) alongside positive control Fusicoccin A (125 μM) against DMSO vehicle treatment of human 14-3-3σ protein (300 nM) and the phosphorylated ERα peptide (20 nM). The top hit, EN171, is shown in red. **(b)** Structure of EN171 with the cysteine-reactive acrylamide moiety highlighted in red. **(c)** EN171 showed an EC50 of 9 μM on 14-3-3σ with ERα, 45 μM with YAP, and 107 μM with TAZ. 14-3-3σ protein (300 nM) and the phosphorylated peptide (20 nM) were incubated with EN171 for 1 h. **(d)** Attenuation of EN171-mediated stabilization between 14-3-3σ and ERa, YAP, or TAZ with the 14-3-3σ inhibitor R18. 14-3-3σ protein (300 nM), the phosphorylated peptide (20 nM), and 14-3-3σ inhibitor R18 (100 μM) were incubated with EN171 for 4 h (ERα) or 1 h (TAZ and YAP). Data shown in **(c)** and **(d)** are average ± sem from n=3 biologically independent replicates per group.

We next tested EN171 against eight additional substrates to better understand the selectivity of EN171 stabilization across known 14-3-3 substrates including c-MYC, USP8, p65, c-RAF, TP53, CDC25B, SOS1, and ABL1 **(Figure S1)**. EN171 did not stabilize 14-3-3 interactions with phosphopeptide substrates from USP8, p65, c-RAF, TP53, SOS1, or ABL1 at 1 mM, but did stabilize interactions with MYC and CDC25B with EC_50_s of 253 and 187 μM, respectively **(Figure S1)**.

### Characterization of 14-3-3 Covalent Stabilizer EN171

We next sought to determine the site of covalent interaction of EN171 with 14-3-3. Tandem mass spectrometry (MS/MS) analysis of tryptic digests from 14-3-3σ incubated with EN171 surprisingly revealed two sites of modification—the expected C38 as well as C96 **(Figure 2a)**. To further characterize EN171 interactions with 14-3-3σ, we synthesized an alkyne-functionalized analog of EN171, EN171-alkyne **(Figure 2b)**. EN171-alkyne dose-responsively and covalently labeled pure 14-3-3σ protein by gel-based activity-based protein profiling (ABPP) approaches **(Figure 2c)** ^13,14^. We demonstrated that neither C38S or C96S mutations in 14-3-3σ prevented EN171-alkyne labeling, but mutation of both C38 and C96 to serine completely ablated probe labeling **(Figure 2d)**. Mutation of either C38 or C96 to serine also significantly attenuated EN171-mediated stabilization of 14-3-3σ interaction with phosphorylated ERα and TAZ peptide substrates **(Figure 2e)**.

**Figure 2.**
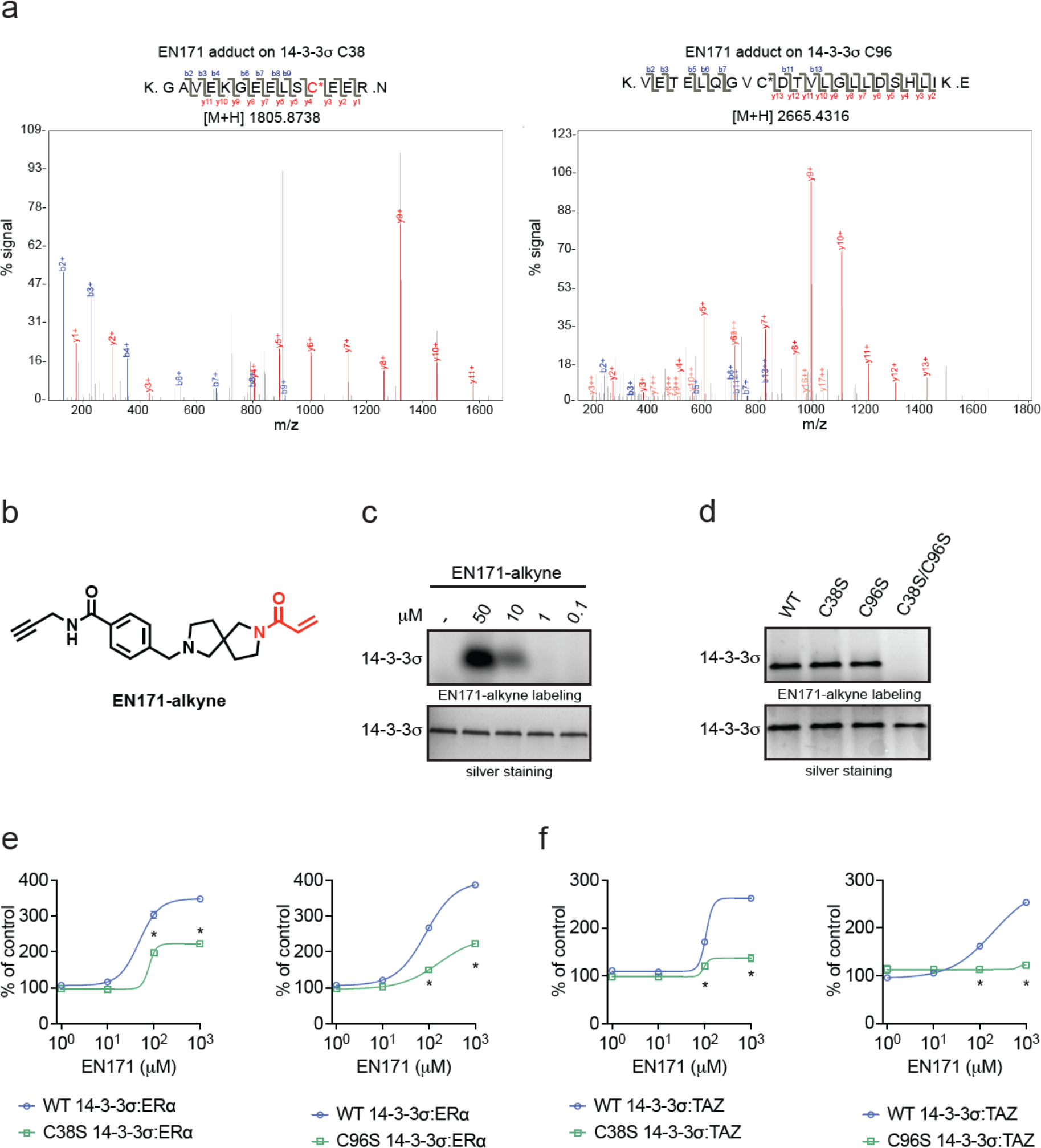
Characterization of 14-3-3 Covalent Stabilizer EN171. **(a)** Mass spectrometry analysis of EN171 modification on 14-3-3σ. Pure human protein 14-3-3σ was incubated with EN171 (50 μM) for 30 min and tryptic digests from the protein were subsequently analyzed for the EN171 covalent adduct on a cysteine. Shown is the MS/MS data for EN171 modification on 14-3-3σ C38 and C96. **(b)** Structure of the alkyne-functionalized analog of EN171 (EN171-alkyne). **(c)** Gel-based ABPP showing that EN171-alkyne labeling of pure human 14-3-3σ protein. 14-3-3σ was incubated with DMSO vehicle or EN171-alkyne for 1 hour. Rhodamine-azide was appended to probe-modified protein by CuAAC, subjected to SDS/PAGE, and analyzed by in-gel fluorescence. Protein loading was assessed by silver staining. **(d)** Gel-based ABPP showing that mutation of both C38 and C96 to serines completely ablated EN171-alkyne probe labeling. 14-3-3σ WT or mutant protein were incubated with EN171-alkyne (50 μM) for 1 hour. **(e)** FP assay showing that mutation of either C38 or C96 to serines also significantly attenuated EN171-mediated stabilization of 14-3-3σ interaction with phosphorylated ERα and TAZ peptide substrates. 14-3-3σ WT or mutations (300 nM) and the phosphorylated peptide (20 nM) were incubated with EN171 for 4 hours. Data shown in **(c-f)** are from n=3 biologically independent replicates per group. Data shown in **(e**,**f)** are average ± sem and significance is shown as *p<0.05 compared to WT 14-3-3 treatment groups.

### Cellular Activity of EN171

Using the EN171-alkyne probe, we next assessed target engagement of this probe in cells. We treated cells with the EN171-alkyne probe and subsequently appended a biotin enrichment handle through copper-catalyzed azide-alkyne cycloaddition (CuAAC), pulled down probe-modified proteins, and demonstrated enrichment of 14-3-3σ, but not unrelated proteins such as GAPDH **(Figure 3a)**. To assess the overall selectivity of EN171, we performed a EN171-alkyne pulldown quantitative proteomic experiment identifying proteins that were outcompeted by excess EN171 pre-treatment. Out of 5266 proteins quantified, we identified 65 proteins, including 14-3-3σ that were significantly outcompeted by EN171 by over 2-fold **(Figure 3b; Table S2)**. To further assess intracellular target engagement and overall selectivity, we performed cysteine chemoproteomic to map the proteome-wide cysteine-reactivity of EN171 in cells using mass spectrometry-based ABPP ^15–17^. We observed significant, albeit low degree of engagement of both C38 and C96 of 14-3-3σ in cells with control versus treated ratios of 1.2 and 1.14, respectively, indicating 12-17 % engagement **(Figure S2; Table S3)**. With regards to overall cysteine-reactivity, we identified 46 targets that were significantly engaged by >75 % (ratio of 4) out of >15,000 probe-modified cysteines quantified **(Figure S2; Table S3)**. Overall, our data indicated that we were engaging 14-3-3σ with a moderate degree of selectivity in complex proteomes or in cells. Among the off-targets of EN171, we did not observe obvious targets that would modulate transcription factor signaling and surmised that we could distinguish 14-3-3-mediated effects through appropriate genetic validation studies. We next assessed whether EN171 impaired ERα and YAP and TAZ-mediated Hippo transcriptional activity in cells using luciferase reporter assays. EN171 dose-responsively impaired both ERα and Hippo transcriptional activity in T47D and MCF7 breast cancer cells, respectively **(Figure 3c-3d)**. These inhibitory effects on ERα and Hippo transcriptional activity were significantly attenuated upon knockdown of 14-3-3σ, indicating that these transcriptional inhibitory effects were on-target **(Figure 3e-3h)**.

**Figure 3.**
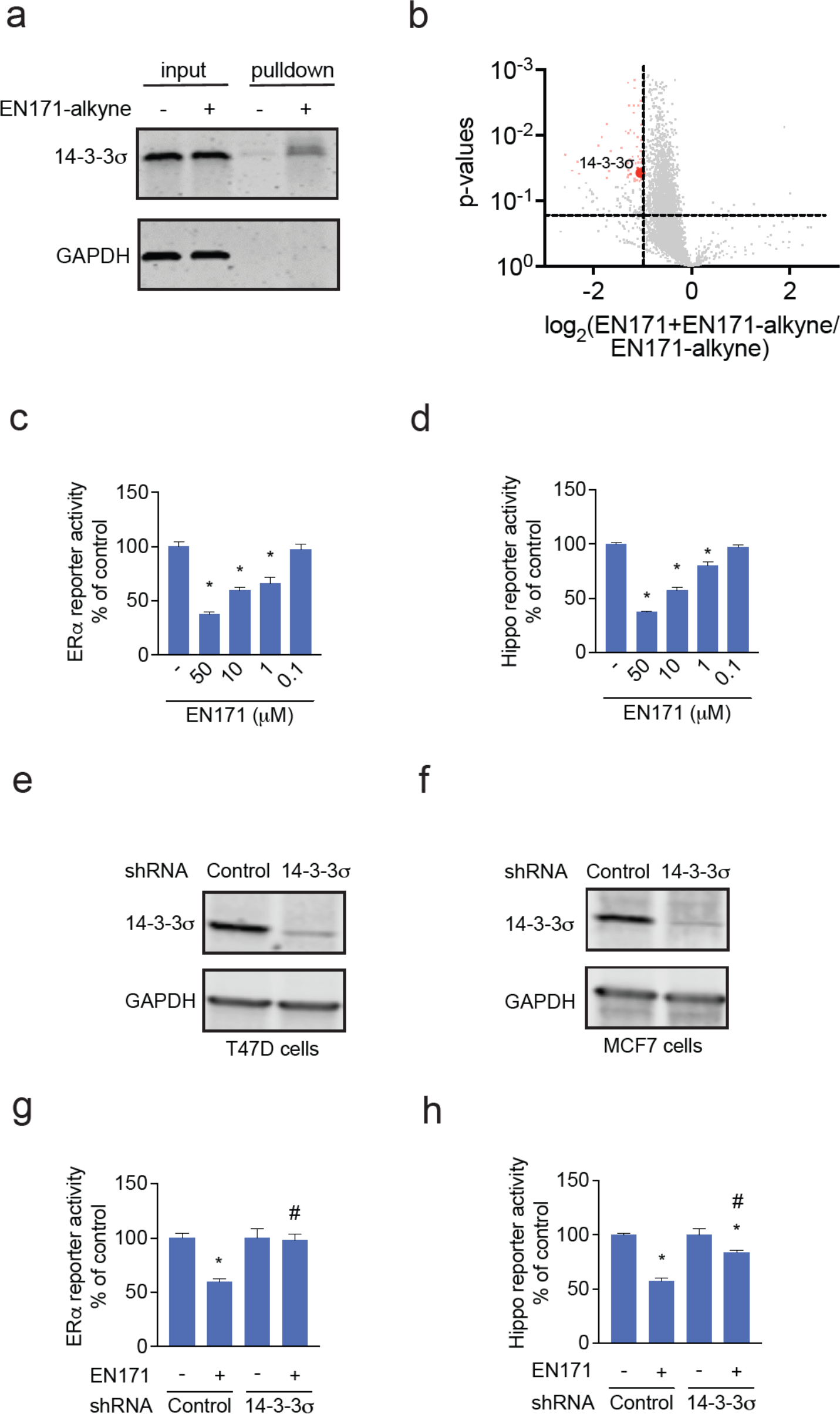
Cellular Activity of EN171. **(a)** Pulldown of endogenous 14-3-3σ in T47D cells using EN171-alkyne. T47D cells were treated with DMSO vehicle or EN171-alkyne probe (50 μM) for 1 h. An azide-functionalized biotin handle was appended onto probe-labeled proteins by CuAAC and these proteins were subsequently streptavidin-enriched. Resulting pulled down proteins were analyzed by SDS–PAGE and Western blotting for 14-3-3σ and loading control GAPDH. **(b)** EN171-competed targets from EN171-alkyne pulldown proteomics. T47D cell lysate were spiked with 14-3-3σ pure protein, pre-treated with DMSO vehicle or EN171 (200 μM) for 1 h at 25 °C prior to EN171-alkyne probe labeling (40 μM) at 4 °C overnight. Probe-modified proteins were subjected to CuAAC with an azide-functionalized biotin and enriched by streptavidin after which eluates were analyzed by TMT-based quantitative proteomics. **(c-d)** Luciferase Reporter Assays showing that EN171 dose-responsively impaired ERα and YAP/TAZ-mediated Hippo transcriptional activity in T47D and MCF7 breast cancer cells, respectively. Cells were treated with DMSO vehicle or EN171 at 37°C for 24h before proliferation was assayed by Hoechst staining or WST-1 and luciferase signal was measured by Steady-Glo^®^ Luciferase Assay System. **(e-f)** 14-3-3σ knockdown with shRNA in T47D and MCF7 cells. Western blot for 14-3-3σ and loading control GAPDH in transfected cells. **(g-h)** Attenuation of ERα and Hippo transcriptional activity by knockdown of 14-3-3σ. shControl and sh14-3-3σ T47D and MCF7 cells were treated with DMSO vehicle or EN171 (10 μM) for 24h. Data shown in **(a-h)** are from n=3 biologically independent replicates per group. Data in **(c**,**d**,**g**,**h)** are average ± sem. Significance is expressed as *p<0.05 compared to vehicle-treated controls, and #p<0.05 compared to EN171 treated shControl cells in **(g**,**h)**.

### Using EN171 as a Covalent Recruiter for 14-3-3σ to Sequester Neo-Substrates

Thus far, chemical approaches to target 14-3-3 proteins have focused on stabilizing interactions between 14-3-3 and native phosphorylated substrates ^1,2^. However, whether 14-3-3 targeting small-molecules can be used to sequester nuclear neo-substrates that are not native 14-3-3 substrates into the cytosol have not been investigated. To enable targeted protein localization of potential nuclear neo-substrates into the cytosol using 14-3-3, we exploited EN171 as a covalent recruiter for 14-3-3σ in a heterobifunctional molecule linking it to the BET family inhibitor JQ1 to sequester nuclear BRD4 into the cytosol to synthesize QS57 **(Figure 4a)**. QS57 treatment in both HEK293T cells and T47D breast cancer cells led to reduced nuclear BRD4 protein levels, and substantially increased cytosolic BRD4 **(Figure 4b-4c)**. To confirm that this cytosolic BRD4 sequestration was mediated through 14-3-3, we first knocked down 14-3-3 using short hairpin RNA (shRNA). Partial knockdown of 14-3-3σ modestly attenuated BRD4 cytosolic sequestration **(Figure 4d)**. Given previous studies with Proteolysis Targeting Chimeras using covalent E3 ligase recruiters demonstrating that only minimal engagement of E3 ligases may be required to achieve robust degradation of their targets, we postulated that we may require full knockout of 14-3-3 to achieve more robust attenuation of BRD4 sequestration. Consistent with this premise, knockout of 14-3-3σ led to complete attenuation of QS57-mediated BRD4 cytosolic sequestration in HCT116 colorectal cancer cells **(Figure 4e)**.

**Figure 4.**
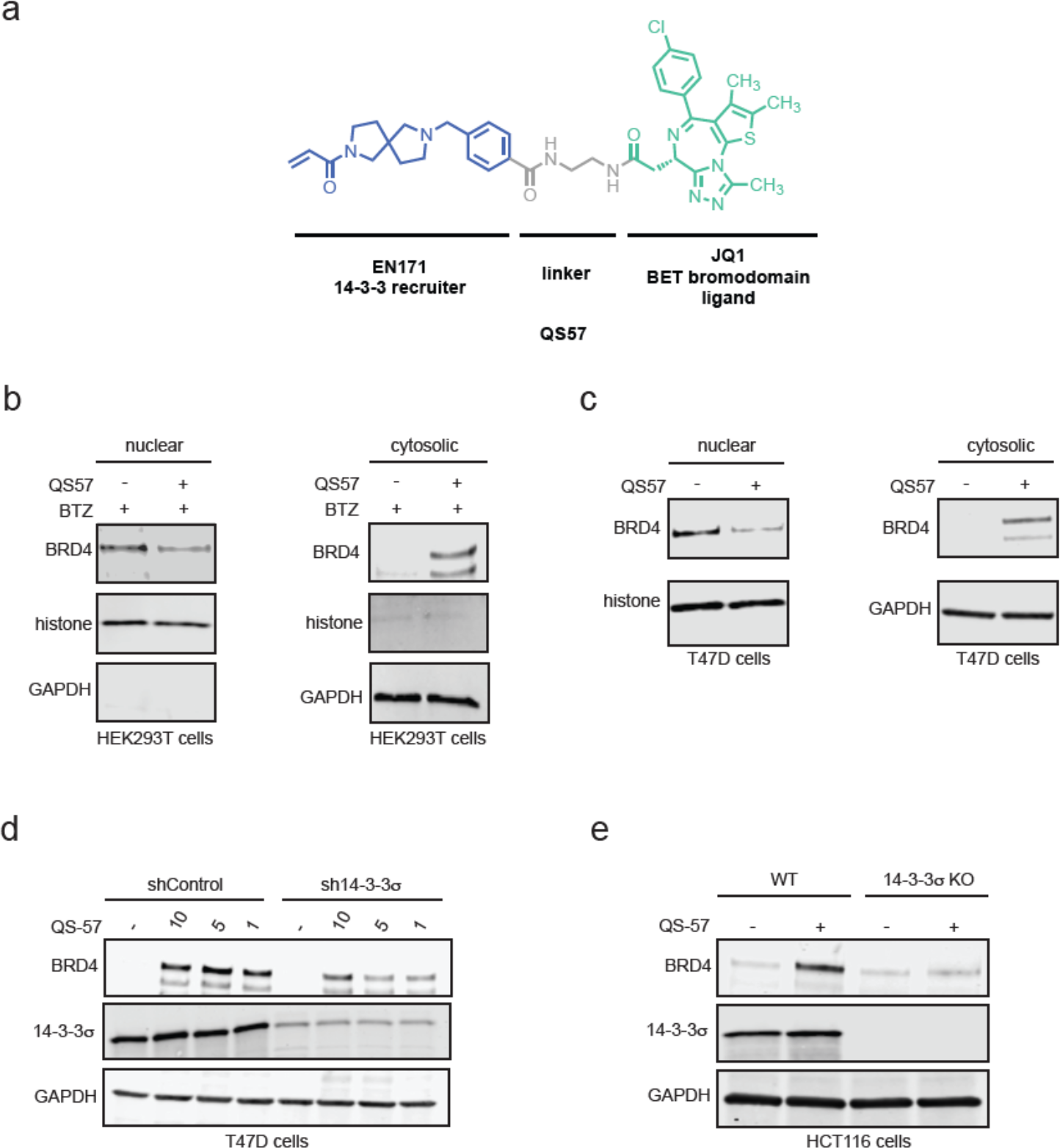
Heterobifunctional Recruiter for 14-3-3σ to Sequester Neo-Substrates. **(a)** Structures of QS57 using covalent 14-3-3σ recruiter in blue linked via a C2 alkyl linker in grey to the BET family inhibitor JQ1 in green. **(b-c)** Sequestration of neo-substrate proteins BRD4 from the nucleus into the cytosol. HEK293T cells were pre-incubated with bortezomib (BTZ) (1 μM) for 1h, prior to treatment of DMSO vehicle or QS57 (100 μM) for 1 d. T47D cells were treated with DMSO vehicle or QS57 (25 μM) for 1 d. **(d)** Attenuation of BRD4 cytosolic sequestration by sh14-3-3σ knockdown. shControl and sh14-3-3σ T47D cells were treated with DMSO vehicle or QS57 (1-10 μM) for 24 h. **(e)** Attenuation of BRD4 cytosolic sequestration by 14-3-3σ knockout. WT and 14-3-3σ knockout HCT116 cells were treated with DMSO vehicle or QS57 (100 μM) for 24h. Data shown in **(d)** are from n=2 biologically independent replicates per group. Data shown in **(b**,**c**,**e)** are from n=3 biologically independent replicates per group. BRD4, 14-3-3σ, and loading control cytosolic GAPDH or nuclear histones were assessed by Western blotting.

## Conclusion

Overall, our study uncovers a covalent molecular glue EN171 that targets 14-3-3σ to stabilize interactions between not only previously stabilized substrates such as ERα, but also less tractable transcription factor substrates such as YAP and TAZ, leading to impaired ERα and Hippo transcriptional activity. We also demonstrate that this covalent 14-3-3 molecular glue can also be used as a covalent 14-3-3σ recruiter to sequester neo-substrates that are not native 14-3-3 substrates such as BRD4 out of the nucleus and into the cytosol. While EN171 still requires significant medicinal chemistry efforts to improve potency, selectivity, and drug-like properties, our findings demonstrate the utility of using covalent ligand libraries to discover novel molecular glues for proteins beyond degradation. Our work expands upon substantial literature precedent from other research groups in using either natural product scaffolds, fully synthetic small-molecule libraries, or disulfide tethering approaches to identify stabilizers of 14-3-3 and its native phosphorylated substrates ^1–12^. Interestingly, EN171 appears to act through targeting two cysteines that are not spatially close to each other— C38 and C96. We still do not understand how these two cysteines work together to exert the actions of EN171 and whether there is potentially an order of reactivity, where EN171 reacts with C38 first and then reveals a secondary site of labeling. Future structural biology studies may reveal the mechanism of EN171 binding to 14-3-3.

We also demonstrate here that covalent 14-3-3 recruiters can also be used in a heterobifunctional mode to sequester neo-substrate proteins from the nucleus and into the cytosol with a fully synthetic and non-genetically encoded approach. This further builds on recent work where genetically encoded systems were developed to sequester cytosolic proteins into the nucleus ^18,19^. These types of targeted protein localization approaches could be a new therapeutic modality for disrupting or enhancing the functions of proteins by localizing or sequestering them out of their native functional contexts. Taken more broadly, our studies demonstrate the utility of covalent ligand screening approaches for discovering novel molecular glues and therapeutic modalities.

## Acknowledgement

We thank the members of the Nomura Research Group for critical reading of the manuscript. This work was also supported by Apple Tree Partners, the Mark Foundation for Cancer Research ASPIRE Award and the National Institutes of Health (R35CA263814). We also thank Drs. Hasan Celik, Alicia Lund, and UC Berkeley’s NMR facility in the College of Chemistry (CoC-NMR) for spectroscopic assistance. Instruments in the College of Chemistry NMR facility are supported in part by NIH S10OD024998.

## Author Contributions

QS, DKN conceived of the project idea, designed experiments, performed experiments, analyzed and interpreted the data, and wrote the paper. QS, TNG, IP, DKN performed experiments, analyzed and interpreted data, and provided intellectual contributions.

## Methods

### Materials

Cysteine-reactive covalent ligand libraries purchased from Enamine and also from internally sourced compounds that were previously described ^14,17,20–23^. EN171 was purchased from Enamine (catalog number EN300-7530357).

### Fluorescence Polarization Assay

Fluorescein-labeled peptides (20 nM), 14-3-3 protein (300 nM), and ligands (50 mM stock solution in DMSO) were diluted in buffer (10 mM HEPES pH 7.5, 150 mM NaCl, 0.1% TWEEN-20). Final DMSO concentration in the assay was always 1%. Dilution series of 14-3-3 protein or fragments were made in black, round-bottom 384-microwell plates (Corning) in a final sample volume of 10 μL in triplicates. Fluorescence anisotropy measurements were performed directly and after 1 hour incubation at room-temperature, using a Tecan plate reader (filter set λ_ex_: 485 ± 20 nm, λ_em_: 535 ± 25 nm). Data reported are at end point. EC_50_ values were obtained from fitting the data in GraphPad Prism.

### LC/MS-MS Analysis of EN171 Interactions with 14-3-3σ

14-3-3σ 10 µg in 100 μL PBS was incubated for 30 min at rt with EN171 (50 μM). The sample was precipitated by addition of 10 µL of 100% (w/v) TCA and cooled to -80 °C for 1 h. The sample was then spun at max speed for 10 min at 4°C and washed three times with ice cold 0.01 M HCl/90% acetone solution. The pellet was resuspended in 4 M urea containing 0.1 % Protease Max (Promega Corp. V2071) and diluted in 40 mM ammonium bicarbonate buffer. Next, the sample was reduced with 10 µl of 10 mM TCEP and incubated at 60 °C for 30 min. Afterwards, 10 µl of 150mM iodoacetamide was added and incubated at room temperature for 30 minutes. The sample was then diluted 50% with PBS before addition of sequencing grade trypsin (1 μg per sample, Promega Corp, V5111) and incubated overnight at 37 °C. Samples were acidified to a final concentration of 5 % formic acid and centrifuged at 13,200 rpm for 30 min. The supernatant was transferred to a new tube and fractionated using high pH reversed-phase peptide fractionation kits (ThermoFisher, 84688) following manufacturer’s recommendation.

### Gel-based ABPP

Gel-based ABPP methods were performed as previously described ^[14]^. Recombinant pure human 14-3-3σ (Origene) or mutations ^[12]^ (0.5 µg), was treated with DMSO or EN171-alkyne probe (0.1-50 µM final concentration) for 1 hour at rt in a volume of 50µL of PBS with or without 1 mg/mL BSA. CuAAC was performed to append rhodamine-azide (20 µM final concentration) onto alkyne probe-labeled proteins. Samples were then diluted with 30 μL of 4 x reducing Laemmli SDS sample loading buffer (Alfa Aesar) and heated at 95 °C for 5 min. The samples were separated on precast 4-20% TGX gels (Bio-Rad Laboratories, Inc.). The gel was analyzed by in-gel fluorescent signal using a ChemiDoc MP (Bio-Rad). Imaged gels were stained using Pierce™ Silver Stain Kit (Thermo Scientific™, 24612) following manufacturer’s instruction.

### Cell Culture

Estrogen Receptor Luciferase Reporter T47D Stable Cell Line (obtained from Signosis, SL-0002) was cultured in RPMI-1640 containing 10% (v/v) fetal bovine serum (FBS) and 1% Penicillin/Streptomycin. During experiments, the cells were grown in phenol red-free DMEM supplemented with 10% charcoal-treated FBS (HyClone) and 1 nM β-Estradiol (E2). Hippo Pathway/TEAD Luciferase Reporter MCF7 Cell Line (obtained from BPS Bioscience, 60618) was cultured in MEM medium supplemented with 10% FBS, 1% non-essential amino acids, 1 mM Na pyruvate, 1% Penicillin/Streptomycin plus 400 µg/ml Geneticin and 10 µg/ml insulin. HEK293T cell line was sourced from the UC Berkeley Cell Culture Facility and cultured in DMEM containing 10% (v/v) fetal bovine serum (FBS) and 1% Penicillin/Streptomycin. HCT116 SFN knockout cell line (obtained from Horizon, HD R02-001) was cultured in RPMI-1640 containing 10% (v/v) fetal bovine serum (FBS) and 1% Penicillin/Streptomycin. All the cell lines were maintained at 37°C with 5% CO_2_.

### Luciferase Reporter Assays

Cells were seeded in a 96 well plate at 10,000/well in a 100 µl volume and allowed to adhere overnight. Ligands or DMSO vehicle were added to the wells. Cells were incubated at 37°C for 24 h before proliferation was assayed by Hoechst staining or WST-1 and luciferase signal was measured by Steady-Glo^®^ Luciferase Assay System (Promega, E2520).

### SFN Lentiviral Knockdown Studies

For each replicate, two 1.5 mL tubes were used. To the first tube, 2 µg of following lentiviral plasmids – shRNA construct of SFN, psPAX2 (carries GAG, REV, polgenes) and pMD2G (carries VSVG pseudotyping gene) – were dissolved in 1.2 mL of Gibco™ Opti-MEM™ I Reduced Serum Medium (catalogue no. 31985-062). To the second tube, lipofectamine™ 2000 (Invitrogen™, 11668019) was diluted in 1.2 mL of Gibco™ Opti-MEM™ I Reduced Serum Medium. Each tube was sat undisturbed for 5 minutes, and the two were mixed into one. After 30 minutes of incubation at a room temperature without mixing, each combined tube was added to HEK293T cells at 40% confluency. Media was replaced in following day, and the cells were incubated for 48 hours. At the infection day, the incubated viral soup was collected from HEK293T cells. After a round of filtration using 0.45 µm filter, the viral soup was combined with equal volume of target cell line’s media, and polybrene (Sigma-Adrich, TR-1003-G) was added to the combined media in 1:1000 dilution. The lentiviral mixture was then added to the target cells, and the media was replaced with fresh media in the following day. After 48 hours, infected cells were selected using puromycin.

### Pulldown of 14-3-3σ Using EN171-Alkyne

T47D cells at 80% confluency were treated with DMSO or EN171-Alkyne (50 mM). After 1 hours of incubation, cells were harvested and lysed. Each lysate was normalized to a concertation of 5 mg/mL, and 500 µL of each lysate was transferred to separate tube. To each tube containing 500 µL of cell lysate, 10 µL of 10 mM biotin picolyl azide (Sigma Aldrich, 900912) in DMSO, 10 µL of 50 mM TCEP in H_2_O, 10 µL of 50 mM CuSO_4_ in H_2_O, and 30 µL of TBTA ligand (1.3 mg/mL in 1:4 DMSO/*t*BuOH, Cayman chemical, 18816) were added. The reaction mixture was incubated at 23 °C for 60 minutes, and the reaction was quenched by proteins precipitation. After washing the protein pellets three times with cold MeOH (4 °C), the pellets were redissolved in 200 µL of 1.2 % SDS/PBS (w/v). After heating the samples at 90 °C for 5 minutes, 10 µL of each sample was set aside for the western blot analysis (input control). 1 mL of PBS was added to the remaining sample to reduce the total SDS concentration to less than 0.2 % SDS/PBS (w/v). 100 µL of streptavidin agarose beads (ThermoFisher, 20353) were added to the lysates containing tubes and the samples were incubated at 4 °C on a rotator for overnight. After incubation, samples were brought to 23 °C, and beads were spined down in a centrifuge (1400 g, 3 min). The supernatant was removed, and beads were washed three more times with 500 µL of PBS and 500 µL of H_2_O. The beads were then suspended in 30 µL of Laemmli SDS sample loading buffer and heated to 95 °C for 10 minutes. Along with the input control, the resulting samples were subjected to western blot analysis.

### EN171-competed targets from EN171-alkyne pulldown proteomics

T47D cells were harvested, lysed, and the proteome concentration was adjusted to 5 mg/mL in 500 μL of PBS using the BCA assay. T47D cell lysate were spiked with 14-3-3σ pure protein (0.5 μM), pre-treated with DMSO vehicle or EN171 (200 μM) for 1 h at 23 °C prior to EN171-alkyne probe labeling (40 μM) at 4 °C overnight. To each tube containing cell lysate, the following reagents were added: 10 μL of 10 mM biotin picolyl azide (Sigma Aldrich, 900912) in DMSO, 10 μL of 50 mM TCEP in H_2_O, 10 μL of 50 mM CuSO_4_ in H_2_O, and 30 μL of TBTA ligand (1.7 mM in 1:4 DMSO/tBuOH, Cayman Chemical, 18816). The reaction mixture was incubated at 23 °C for 60 minutes, and the reaction was quenched by protein precipitation. Precipitated pellets were washed using 500 μL of MeOH and centrifuged again to yield white pellets. Samples were resuspended in 1.2% SDS-PBS (1 mL), completely dissolved, and heated to 90 °C for 5 minutes. The soluble proteome was then diluted with 5 mL of PBS and further incubated with high-capacity streptavidin-agarose beads (100 μL/sample, ThermoFisher Scientific, 20357). Beads and lysates were incubated overnight at 4 °C with rotation. On the following day, beads were suspended and washed three times with 0.1% SDS-PBS, PBS, and H2O. Washed beads were resuspended in 6 M Urea/PBS (500 μL), and the samples were further treated with DTT and iodoacetamide. After removing the supernatant, beads were resuspended in 100 μL of 50 mM TEAB and enzymatically digested overnight using sequencing-grade trypsin (Promega, V5111). Digested peptides were eluted through centrifugation and labeled using commercially available TMTsixplex tags (ThermoFisher, P/N 90061). After labeling, 35 μg of each labeled sample was combined and dried using a vacufuge. Dried samples were redissolved with 300 μL of 0.1% TFA in H_2_O and further fractionated using high-pH reversed-phase peptide fractionation kits (ThermoFisher, P/N 84868) following the manufacturer’s protocol. Dried fractions were then resuspended in 25 μL of 0.1 % Formic acid/H_2_O (w/v) to be analyzed by LC-MS/MS.

### Mass Spectrometry Analysis

Mass spectrometry analysis was performed on an Orbitrap Eclipse Tribrid Mass Spectrometer with a High Field Asymmetric Waveform Ion Mobility (FAIMS Pro) Interface (Thermo Scientific) with an UltiMate 3000 Nano Flow Rapid Separation LCnano System (Thermo Scientific). Off-line fractionated samples (5 μl aliquot of 15 μl sample) were injected via an autosampler (Thermo Scientific) onto a 5 μl sample loop which was subsequently eluted onto an Acclaim PepMap 100 C18 HPLC column (75 μm x 50 cm, nanoViper). Peptides were separated at a flow rate of 0.3 μl/min using the following gradient: 2 % buffer B (100 % acetonitrile with 0.1 % formic acid) in buffer A (95:5 water:acetonitrile, 0.1 % formic acid) for 5 min, followed by a gradient from 2 to 40 % buffer B from 5 to 159 min, 40 to 95 % buffer B from 159 to 160 minutes, holding at 95 % B from 160-179 min, 95 % to 2 % buffer B from 179 to 180 min, and then 2 % buffer B from 180 to 200 min. Voltage applied to the nano-LC electrospray ionization source was 2.1 kV. Data was acquired through an MS1 master scan (Orbitrap analysis, resolution 120,000, 400-1800 m/z, RF lens 30 %, heated capillary temperature 250 °C) with dynamic exclusion enabled (repeat count 1, duration 60 s). Data-dependent data acquisition comprised a full MS1 scan followed by sequential MS2 scans based on 2 s cycle times. FAIMS compensation voltages (CV) of -35, -45, and -55 were applied. MS2 analysis consisted of quadrupole isolation window of 0.7 m/z of precursor ion followed by higher energy collision dissociation (HCD) energy of 38 % with a orbitrap resolution of 50,000.

### Western Blotting

Nuclear and Cytoplasmic extraction were separated using NE-PER™ Nuclear and Cytoplasmic Extraction Reagents (Thermo Fisher, 78833) following manufacturer’s instruction. Proteins were separated by precast 4-20% Criterion TGX gels (Bio-Rad) and transferred to nitrocellulose membrane using the Trans-Blot Turbo transfer system (Bio-Rad). After the transfer, membrane was blocked with Tris-buffered saline containing Tween 20 (TBST) containing 5% bovine serum albumin (BSA) for 1 hours at RT. After the blocking, targeted proteome was probed with primary antibody in TBST with 5% BSA (Primary antibody dilution condition from manufacturer). Incubation with primary antibody was performed overnight in cold room (4 °C), and the primary antibody was washed three times with TBST. Membrane was then incubated in the dark with IR680 or IR800-conjugated secondary antibodies (1: 10000 dilution). After 1 hour incubation at RT, blot was washed 3 times with TBST, and the blots were visualized using Odyssey Li-Cor fluorescent scanner. Following antibodies were used for this study: 14-3-3σ (Santa Cruz, sc-100638; Abcam, ab151504), GAPDH (Proteintech, 60004-1-1G-150L), BRD4 (Abcam, ab128874), Histone (Cell Signaling Technology, 14269S), IRDye 680RD Goat anti-Mouse (LI-COR 926-60870), and IRDye 800CW Goat anti-Rabbit (LICOR 926-32211).

