## Supporting Information for "Covalent 14-3-3 Molecular Glues and Heterobifunctional Molecules Against Nuclear Transcription Factors and Regulators"

**Supporting Table Legends**

**Table S1. Structures of compounds screened. Tab 1** contains the structures of the compounds screened. **Tab 2** contains the fluorescence polarization (FP) screening data in relation to percent of DMSO control of a cysteine-reactive covalent ligand library (125 $\mu$M) screened against human 14-3-3$\sigma$ protein (300 nM) and the phosphorylated ER$\alpha$ peptide (20 nM).

**Table S2. EN171-alkyne pulldown proteomics.** EN171-competed targets from EN171-alkyne pulldown proteomics. T47D cell lysate were spiked with 14-3-3$\sigma$ pure protein, pre-treated with DMSO vehicle or EN171 (200 $\mu$M) for 1 h at 25 $^{o}$C prior to EN171-alkyne probe labeling (40 $\mu$M) at 4 $^{o}$C overnight. Probe-modified proteins were subjected to CuAAC with an azide-functionalized biotin and enriched by streptavidin after which eluates were analyzed by TMT-based quantitative proteomics. Data are from n=3 biologically independent replicates/group.

**Table S3. isoDTB-ABPP of EN171.** isoDTB-ABPP analysis of EN171 in T47D cells. T47D cells were treated with DMSO vehicle or EN171 (100 $\mu$M) for 6 h, after which lysates were labeled with an alkyne-functionalized iodoacetamide probe (IA-alkyne). Control and treated samples were subjected to CuAAC with a desthiobiotin-azide with an isotopically light or heavy handle, respectively. Probe-labeled proteomes were mixed in a 1:1 control/treated ratio, enriched with avidin, digested with trypsin, and eluted with 0.1% formic acid in 50% acetonitrile for isoDTB-ABPP analysis. Probe-modified peptides were analyzed by LC-MS/MS and control vs treated probe-modified peptide ratios were quantified. Data are from n=3 biologically independent replicates/group.

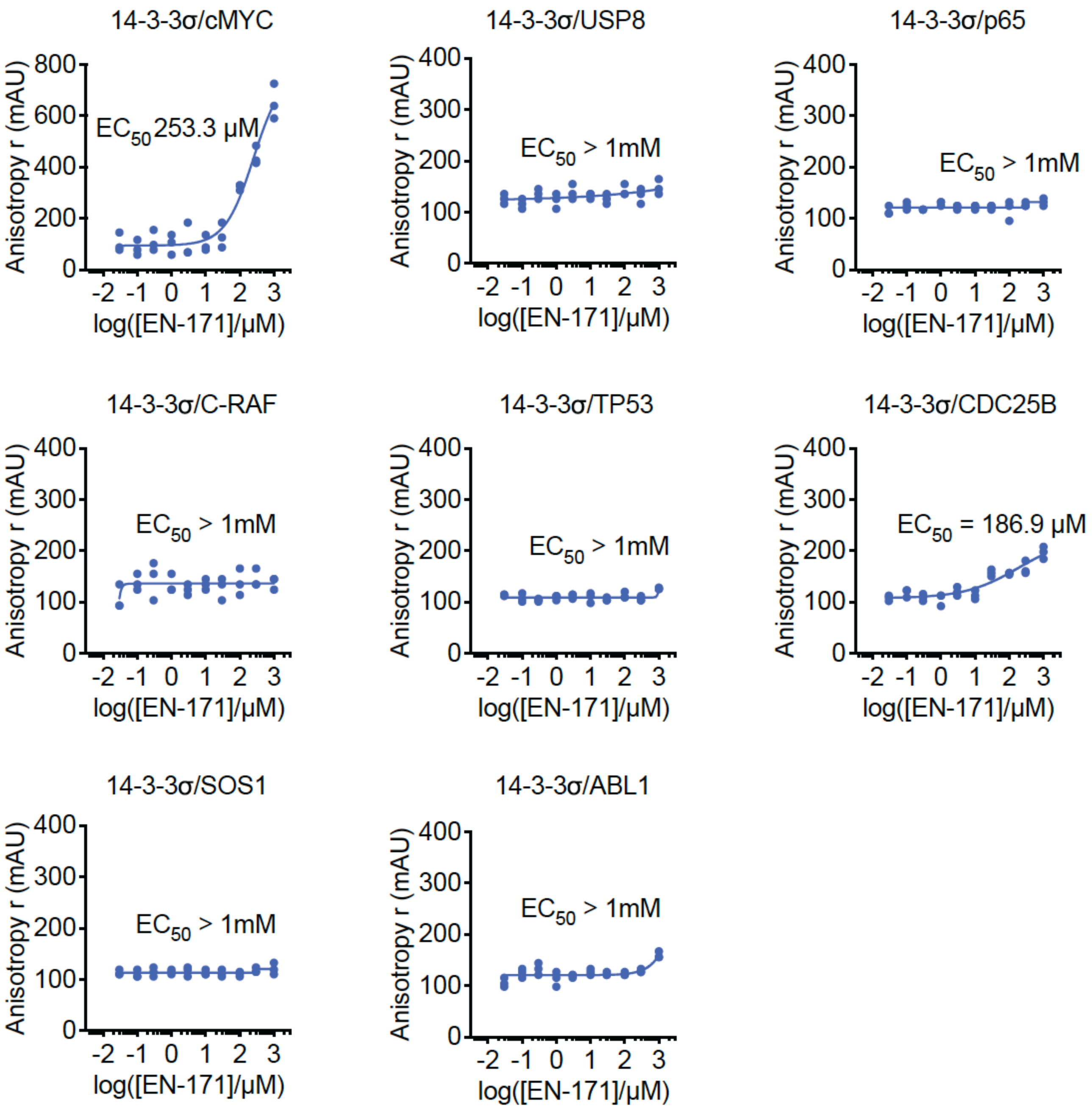


**Figure S1.** Fluorescence polarization (FP) screening of different phosphorylated peptides. 14-3-3σ protein (300 nM) and the phosphorylated peptide (20 nM) were incubated with EN171 for 1 hour at RT. EN171 showed an EC50 of 25.3 μM on 14-3-3σ with c-MYC peptide, 186.9 μM with CDC25B, and did not stabilize 14-3-3 interactions with phosphopeptide substrates from USP8, p65, c-RAF, TP53, SOS1, or ABL1 at 1 mM.

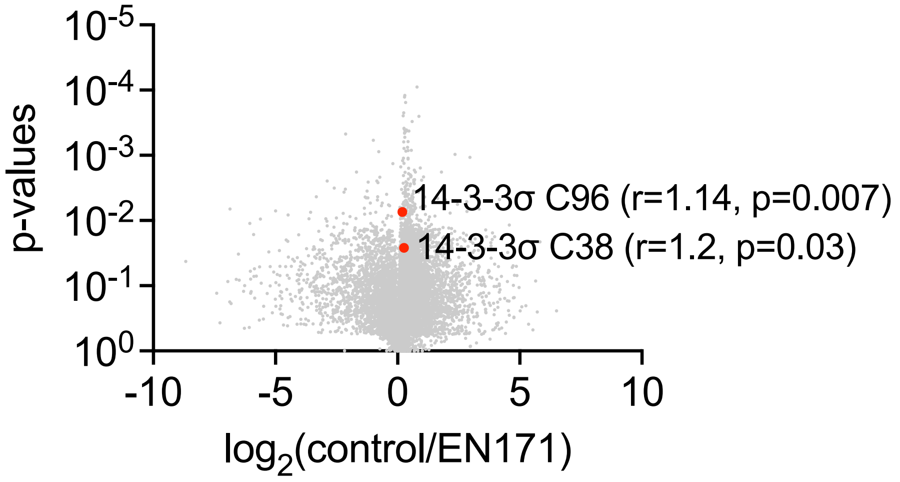


**Figure S2.** isoDTB-ABPP analysis of EN171 in T47D cells. T47D cells were treated with DMSO vehicle or EN171 (100 μM) for 6 h, after which lysates were labeled with an alkyne-functionalized iodoacetamide probe (IA-alkyne). Control and treated samples were subjected to CuAAC with a desthiobiotin-azide with an isotopically light or heavy handle, respectively. Probe-labeled proteomes were mixed in a 1:1 control/treated ratio, enriched with avidin, digested with trypsin, and eluted with 0.1% formic acid in 50% acetonitrile for isoDTB-ABPP analysis. Probe-modified peptides were analyzed by LC-MS/MS and control vs treated probe-modified peptide ratios were quantified. Data are from n=3 biologically independent replicates/group.

### Methods

#### Peptide Sequences

Fluorescein-labeled peptides for FP assay were purchased from GenScript Biotech Corp. Sequences were as follows: *5-FAM*-KYYITGEAEGFPA{pT}V-*COOH* (ER*α*-pp); *5-FAM*-RSH{pS}SPASLQLGT-*CONH2* (TAZ-pp); *5-FAM*-RAH{pS}SPASLQ-*CONH2* (YAP-pp); *5-FAM*-KC{pT}SP-*COOH* (c-Myc-pp); *5-FAM*-KLKRSY{pS}-SPDITQ-*COOH* (USP8-pp); *5-FAM*-EGRSAG{pS}IPGRRS-*COOH* (p65-pp); *5-FAM*-RQRST{pS}TPNVH-*CONH2* (C-Raf-pp); *5-FAM*-SRAHSSHLKSKKGQSTSRHKKLMFK{pT}EGPDSD-*COOH* (TP53); *5-FAM*-QRLFRSP{pS}MPCSVIR-*CONH2* (Cdc25B-pp); *5-FAM*-PRRRPE{pS}APAESS-*COOH* (SOS1-pp); *5-FAM*-GAKDTEWRSV{pT}LPRDLQSTGR-*COOH* (ABL1-pp).

#### Fluorescence Polarization Assay

Fluorescein-labeled peptides (20 nM), 14-3-3 protein (300 nM), and EN171 or other ligands (50 mM stock solution in DMSO) were diluted in buffer (10 mM HEPES pH 7.5, 150 mM NaCl, 0.1% TWEEN-20). Final DMSO concentration in the assay was always 1%. Dilution series of fragments were made in black, round-bottom 384-microwell plates (Corning) in a final sample volume of 10 µL in triplicates. Fluorescence anisotropy measurements were performed directly and after 1 hour incubation at room-temperature, using a Tecan plate reader (filter set $\lambda_{ex}$: 485 ± 20 nm, $\lambda_{em}$: 535 ± 25 nm). Data reported are at end point. $EC_{50}$ values were obtained from fitting the data in GraphPad Prism.

#### IsoDTB-ABPP cysteine chemoproteomic profiling of EN171

T47D cells were treated when they reached 80% confluency, with either DMSO vehicle or 100 µM of EN171 for 6 hours. Cell lysates were prepared using sonication, and proteome concentration was normalized to 2.0 mg/mL. For each biological replicate, two 1 mL aliquots of 2.0 mg/mL were used. Prior to tag labeling, each aliquot was treated with 20 µL of IA-alkyne (200 µM) for 1 hour in 23 °C. Subsequently, each sample set was treated with 120 µL of either heavy or light isoDTB tags containing master mix for 2 hours at 23 °C. Master mix contains 510 µL of TBTA (0.9 mg/mL in 1:4 in DMSO/tBuOH), 165 µL of $CuSO_4$ (12.5 mg/mL in $H_2O$), 165 µL of of TCEP (14.0 mg/mL in $H_2O$), and 160 µL of isoDTB tag (4 mg of either heavy or light tag in DMSO, Click chemistry Tools, 1565). After the reaction, one heavy tagged and one light tagged labeled samples were combined and precipitated in acetone at -20 °C.

After the overnight precipitation, the sample was centrifuged at 7,000 g for 10 minutes in 4 °C. The supernatant was removed, and the precipitated protein samples were resuspended in cold MeOH. Protein pellets were washed with cold MeOH (500 µL, 3 times) and dissolved in 600 µL of 8 M urea in 0.1 M TEAB buffer. Pellets were completely dissolved in the urea solution by sonication, and the urea concentration was adjusted to 2 M urea by adding 1800 μL of 0.1 M TEAB. Two replicates were then combined into a single container, and the combined sample sets were further diluted with 2400 μL of PBS containing 0.2% NP40 (w/v). Proteome was then incubated with high-capacity streptavidin agarose beads (200μL/sample, ThermoFisher, 20357) for overnight at 4 °C on hula mixer.

After incubation, the beads were centrifuged for 1 minute and the supernatant were removed. The remaining beads were then washed in following order: 3 x PBS with 0.1% NP40, 3 x PBS, and 3 x $H_2O$. Next, the protein-bound beads were resuspended in 600 μL of 8 M Urea in 0.1 M of tetraethylammonium bromide (TEAB) buffer, and treated with 30 μL DTT for 45 mins in 37 °C, then 30 μL iodoacetamide for 30 mins in 23 °C, subsequently, 30 μL DTT for 30 mins in 23 °C. After incubation, the beads were centrifuged for 1 minute and the supernatant were removed. The protein-bound beads were resuspended in 400 μL of 2 M Urea in 0.1 M TEAB. 8 μL of 0.5 mg/mL sequencing grade trypsin (Promega, V5111) was then added, and the bead bound proteins were digested overnight at 37 °C. The resulting samples were then diluted with 800 μL of 0.1% NP40 in PBS, and the beads were washed with the same buffer three times and three more times with PBS and $H_2O$. Bead-bound peptides were then eluted with 0.1% formic acid in 50% acetonitrile (v/v). Eluted samples were dried using a vacufuge and re-dissolved in 300 μL of $H_2O$ with 0.1% TFA. Suspended samples were then fractionated using high pH reversed-phase peptide fractionation kits (ThermoFisher, 84688).

Data were extracted in the form of MS1 and MS2 files using Raw Converter (Scripps Research Institute) and searched against the Uniprot human database using ProLuCID search methodology in IP2 v.3-v.5 (Integrated Proteomics Applications, Inc.). Cysteine residues were searched with a static modification for carboxyaminomethylation (+57.02146).

For isoDTB-ABPP, we also searched for up to two differential modifications for methionine oxidation and either the light or heavy isoDTB tags (+561.33872 or +567.34621, respectively). Peptides were required to be fully tryptic peptides. ProLUCID data were filtered through DTASelect to achieve a peptide false-positive rate below 5%. Only those probe-modified peptides that were evident across three out of three biological replicates were interpreted for their isotopic light to heavy ratios. Light versus heavy isotopic probe-modified peptide ratios are calculated by taking the mean of the ratios of each replicate paired light versus heavy precursor abundance for all peptide-spectral matches associated with a peptide. The paired abundances were also used to calculate a paired sample *t*-test *P* value in an effort to estimate constancy in paired abundances and significance in change between treatment and control. *P* values were corrected using the Benjamini–Hochberg method.

**Synthetic Methods and Characterization**

- Starting materials and reagents were purchased from commercial suppliers and used without additional purification step.
- All solvents are reagent grade or HPLC grade.
- Nuclear Magnetic Resonance (NMR) spectra were recorded on Bruker Avance 400, NEO 500, 600 MHz spectrometer. Chemical shift are reported in parts per million (ppm, $\delta$) and calibrated against tetramethylsilane (TMS) or residual solvent signal. $^{1}H$ NMR: $CDCl_3$ (7.26) and DMSO-d6 (2.50) & $^{13}C$ NMR: $CDCl_3$ (77.0) and DMSO-d6 (39.5). Multiplicity is reported as follows: singlet (s), doublet (d), doublet of doublet (dd) doublet of triplet (dt), triplet (t), triplet of doublet (td), quartet (q), and multiplet (m). Coupling constants (J) are reported in Hertz (Hz)
- All reactions were monitored by Thin Layer Chromatography (TLC) from Merck KGaA (TLC Silica gel 60 $F_{254}$). Visualization of TLC was done using UV (254 nm and 365 nm) or chemical stain ($KMnO_4$).
- Purification of product was performed by flash column chromatography using a Biotage Isolera. Elution of compound was monitored with the equipped UV detector, and following Biotage Sfar columns were used based on the scale of synthesis: 5 g, 10 g, or 25 g. If additional purification is needed, compound was further purified using Thermo scientific's semi-prep reversed phase high-performance liquid chromatography equipped (RP-HPLC: Ultimate 3000 HPLC ) equipped with C18 column (Luna® 10 $\mu$m c18(2), 100 Å, Serial #:5293-0084). Elution of sample was monitored using DIONEX UltiMate 3000. Eluting buffer A: 100% distilled water + 0.1% trifluoracetic acid (TFA). Eluting buffer B: 95% acetonitrile + 5% distilled water + 0.1% TFA.
- High-resolution mass spectra (HRMS) were obtained using Q Exactive™ Plus Hybrid Quadrupole-Orbitrap™ Mass Spectrometer.

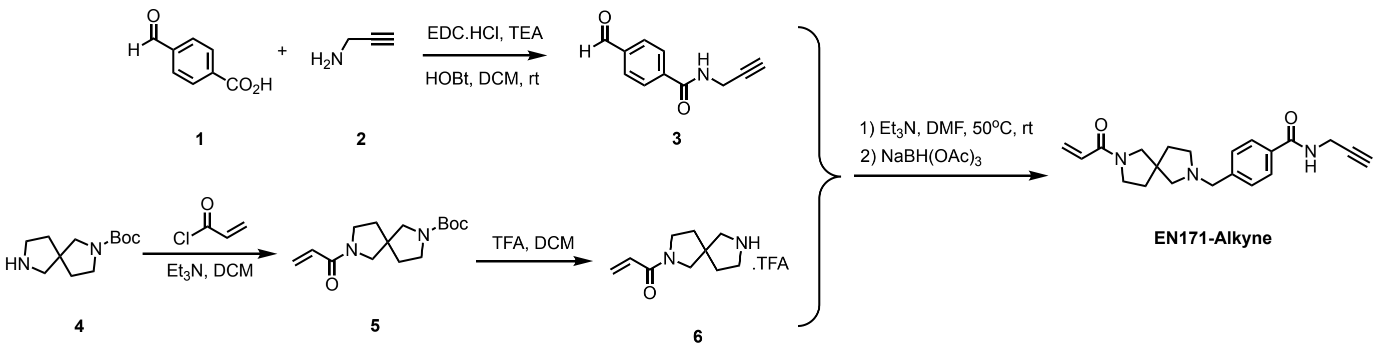


**Procedure for the preparation of EN-171 Alkyne probe:**

**4-formyl-*N*-(prop-2-yn-1-yl)benzamide (3)**

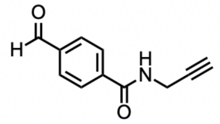

Add the carboxylic acid **1** (216 mg, 1 equiv.), N-(3-Dimethylaminopropyl)-N′-ethylcarbodiimide hydrochloride (320 mg, 1.2 eq), Hydroxybenzotriazole (190 mg, 1 equiv.), and triethylamine (0.2 mL, 1 equiv.) to DCM (3 mL) and stir briefly. Then add alkyne **2** (131 mg, 1 equiv.) and stir overnight at room temperature. The reaction mixture was extracted with DCM, washed with saturated sodium bicarbonate, $NH_4Cl$, and brine (three time each), then dried over $MgSO_4$. After filter and concentration, The product was carried through to the next step without additional purification.

***tert*-butyl 7-acryloyl-2,7-diazaspiro[4.4]nonane-2-carboxylate (5)**

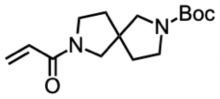


Amine **4** (452 mg, 1 equiv.) was dissolved in DCM (10 mL) with triethylamine (1.4 mL, 5 equiv.). After 10 minutes of vigorous stirring at rt, the reaction was cooled to 0 °C. Acryloyl chloride (0.488 mL, 3 equiv.) was added dropwise to the stirring mixture, then the reaction was warmed to room temperature and stirred overnight. The reaction mixture was concentrated under vacuum and purified through silica gel chromatography (0-80% EtOAc in Hexanes) to result oil **5** (85% yield). $^1$H NMR (400 MHz, $CDCl_3$) δ 6.50 – 6.24 (m, 2H), 5.65 (dd, *J* = 9.7, 3.4 Hz, 1H), 3.72 – 3.54 (m, 2H), 3.43 (td, *J* = 14.0, 7.6 Hz, 4H), 3.30 – 3.11 (m, 2H), 1.98 – 1.78 (m, 4H), 1.41 (s, 9H). $^{13}$C NMR (101 MHz, $CDCl_3$) δ 164.64, 154.48, 128.43, 128.12, 127.87, 79.52, 55.59, 54.96, 54.59, 54.37, 47.31, 46.44, 45.68, 45.05, 44.79, 35.26, 35.11, 34.93, 34.23, 33.50, 28.50. MS (ESI-TOF) Calcd for $C_{10}H_{16}N_2O^+$ [M+H-Boc]$^+$ : 180.13; found: 180.88.

**1-(2,7-diazaspiro[4.4]nonan-2-yl)prop-2-en-1-one (6)**

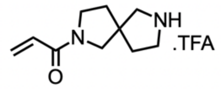


Trifluoroacetic acid (1 mL) was added to a vigorously stirring solution of the Boc protected compound **5** (280 mg) in DCM (3 mL). The reaction mixture was stirred at room temperature for 2 hours. The product mixture was concentrated under vacuum. The product was carried through to the next step without additional purification.

**4-((7-acryloyl-2,7-diazaspiro[4.4]nonan-2-yl)methyl)-*N*-(prop-2-yn-1-yl)benzamide (EN171-Alkyne)**

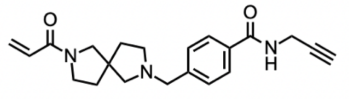

Deprotected amine **6** (55 mg, 1 equiv.) and aldehyde **3** (22 mg, 0.6 equiv.) were dissolved in DMF (2 mL) with triethylamine (0.028 mL, 1 equiv.). And the reaction mixture was stirred at 50 $^oC$ overnight. Subsequently, the reaction mixture was cooled to room temperature and sodium triacetoxyborohydride (85 mg, 2 equiv.) was added in portions. The reaction mixture was stirred at room temperature overnight. The reaction was quenched by adding 5 mL of water. Reactants were extracted with EtOAc five times, and the organic layer was dried over $Na_2SO_4$ and concentrated under vacuum. Resulting oil was purified using flash column chromatography (0-20% MeOH in EtOAc), and white powder was acquired as a product (48% yield). $^1H$ NMR (400 MHz, $CD_3OD$) δ 7.78 (d, *J* = 8.3 Hz, 2H), 7.44 (dd, *J* = 8.4, 1.8 Hz, 2H), 6.54 (ddd, *J* = 16.8, 10.4, 5.0 Hz, 1H), 6.23 (ddd, *J* = 16.8, 3.3, 2.0 Hz, 1H), 5.71 (ddd, *J* = 10.4, 2.0, 0.7 Hz, 1H), 4.13 (d, *J* = 2.5 Hz, 2H), 3.71 (d, *J* = 3.9 Hz, 2H), 3.64 – 3.37 (m, 4H), 2.84 – 2.47 (m, 5H), 2.02 – 1.81 (m, 4H). $^{13}C$ NMR (101 MHz, $CD_3OD$) δ 168.06, 165.34, 142.20, 132.78, 128.72, 128.52, 128.16, 127.05, 79.36, 70.52, 63.11, 63.08, 59.32, 59.30, 57.81, 56.83, 53.25, 53.16, 45.81, 45.00, 37.23, 35.63, 34.90, 34.77, 28.49. HRMS (ESI-TOF) Calcd for $C_{21}H_{26}N_3O_2^+$ $[M+H]^+$ : 352.2020; found: 352.2047.

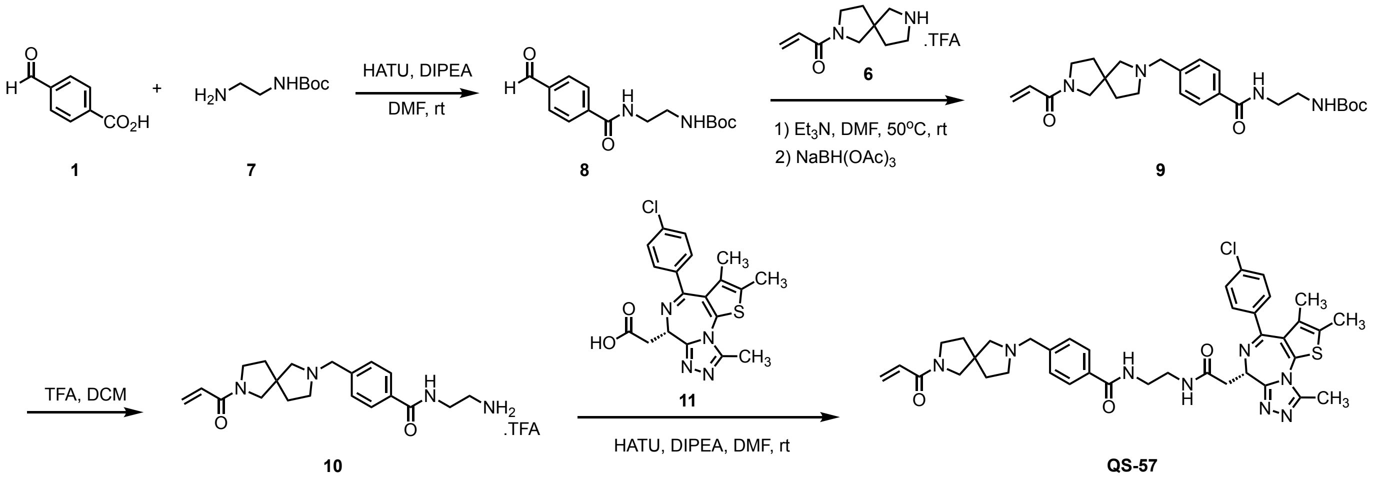

**Procedure for the preparation of QS-57 Bifunctional Molecule:**

***tert*-butyl (2-(4-formylbenzamido)ethyl)carbamate (8)**

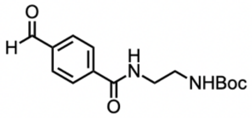


Carboxylic acid **1** (45 mg, 1 equiv.) and hexafluorophosphate azabenzotriazole tetramethyl uranium (152 mg, 1.3 equiv.) was dissolved in DMF (6 mL) along with and N,N-diisopropylethylamine (0.104 mL, 2 equiv.). After stirring at room temperature for 10 minutes, amine **7** (0.063 mL, 1.3 equiv.) was added. The reaction was stirred at room temperature overnight. Organic mixture was extracted using 5 rounds of EtOAc extraction. Combined organic layer was then dried with anhydrous $Na_2SO_4$. After removing solvent in vacuo, compound was purified using flash column chromatography (80-100% EtOAc/Hexane), and white powder was acquired as a product **8** (81% yield). $^1$H NMR (500 MHz, $CDCl_3$) δ 10.00 (s, 1H), 7.92 (d, *J* = 8.1 Hz, 2H), 7.86 (d, *J* = 8.2 Hz, 2H), 7.56 (s, 1H), 5.05 (s, 1H), 3.51 (q, *J* = 5.1 Hz, 2H), 3.36 (q, *J* = 5.9 Hz, 2H), 1.36 (s, 10H). $^{13}$C NMR (126 MHz, $CDCl_3$) δ 191.71, 166.58, 157.95, 139.32, 138.14, 129.78, 127.77, 42.74, 39.72, 28.33. MS (ESI-TOF) Calcd for $C_{10}H_{12}N_2O_2^+$ [M+H-Boc]$^+$ : 192.09; found: 192.89.

***tert*-butyl (2-(4-((7-acryloyl-2,7-diazaspiro[4.4]nonan-2-yl)methyl)benzamido)ethyl)-carbamate (9)**

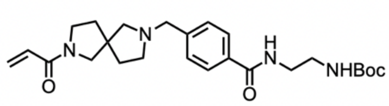


Deprotected amine **6** (55 mg, 1 equiv.) and aldehyde **8** (58 mg, 1 equiv.) were dissolved in DMF (2 mL) with triethylamine (0.028 mL, 1 equiv.). And the reaction mixture was stirred at 50 °C overnight. Subsequently, the reaction mixture was cooled to room temperature and sodium triacetoxyborohydride (85 mg, 2 equiv.) was added in portions. The reaction mixture was stirred at room temperature overnight. Then the reaction was quenched by adding 5 mL of water. Reactants were extracted with EtOAc five times, and the organic layer was dried over $Na_2SO_4$ and concentrated under vacuum. Resulting oil was purified using flash column chromatography (0-20% MeOH in EtOAc), and white powder was acquired as a product **9** (31% yield). $^1$H NMR (500 MHz, $CDCl_3$) δ 7.71 (d, *J* = 7.8 Hz, 2H), 7.32 (t, *J* = 8.7 Hz, 2H), 6.47 – 6.21 (m, 2H), 5.60 (dt, *J* = 9.6, 3.3 Hz, 1H), 5.06 (d, *J* = 9.8 Hz, 1H), 3.66 – 3.31 (m, 10H), 2.60 (dd, *J* = 84.4, 48.7 Hz, 4H), 1.95 – 1.74 (m, 4H), 1.35 (s, 9H). $^{13}$C NMR (126 MHz, $CDCl_3$) δ 167.59, 164.64, 161.12, 128.92, 128.52, 128.11, 127.85, 127.80, 127.67, 127.24, 79.94, 59.56, 53.46, 53.32, 48.69, 46.66, 45.86, 45.23, 40.04, 36.05, 35.21, 28.35. MS (ESI-TOF) Calcd for $C_{25}H_{37}N_4O_4^+$ [M+H]$^+$ : 457.28; found: 457.24.

**4-((7-acryloyl-2,7-diazaspiro[4.4]nonan-2-yl)methyl)-N-(2-aminoethyl)benzamide (10)**

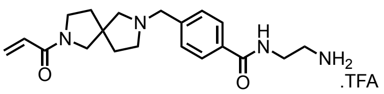


Trifluoroacetic acid (0.2 mL) was added to a vigorously stirring solution of the Boc protected compound **9** (20 mg) in DCM (1 mL). The reaction mixture was stirred at room temperature for 2 hours. The product mixture was concentrated under vacuum. The product was carried through to the next step without additional purification.

**4-((7-acryloyl-2,7-diazaspiro[4.4]nonan-2-yl)methyl)-*N*-(2-(2-((*S*)-4-(4-chlorophenyl)-2,3,9-trimethyl-6*H*-thieno[3,2-*f*][1,2,4]triazolo[4,3-*a*][1,4]diazepin-6-yl)acetamido)ethyl)-benzamide (QS-57)**

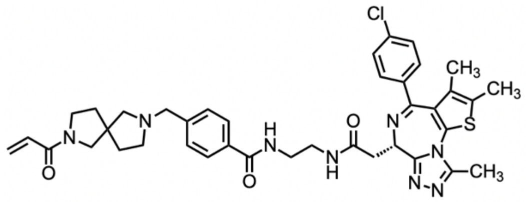


JQ1-acid **11** (40 mg, 2 equiv.) and hexafluorophosphate azabenzotriazole tetramethyl uranium (38 mg, 2 equiv.) was dissolved in DMF (1.5 mL) along with and N,N-diisopropylethylamine (0.026 mL, 6 equiv.). After stirring at room temperature for 10 minutes, amine **7** (20 mg, 1 equiv.) was added. The reaction was stirred at room temperature overnight. Organic mixture was extracted using 5 rounds of EtOAc extraction. Combined organic layer was then dried with anhydrous $Na_2SO_4$. After removing solvent in vacuo, compound was purified using flash column chromatography (80-100% EtOAc/Hexane followed by 2-15% MeOH/DCM), and white powder **QS-57** was acquired as a product (67% yield). $^1$H NMR (500 MHz, MeOD) δ 7.70 (d, *J* = 8.0 Hz, 2H), 7.33 (dd, *J* = 8.4, 2.2 Hz, 2H), 7.31 – 7.17 (m, 4H), 6.54 – 6.39 (m, 1H), 6.21 – 6.04 (m, 1H), 5.68 – 5.57 (m, 1H), 4.54 (dd, *J* = 8.6, 5.8 Hz, 1H), 3.71 – 3.24 (m, 13H), 2.73 – 2.43 (m, 7H), 2.34 (s, 3H), 1.96 – 1.77 (m, 4H), 1.55 (s, 3H). $^{13}$C NMR (126 MHz, MeOD) δ 172.03, 168.70, 165.33, 165.00, 155.60, 150.77, 136.66, 132.16, 130.67, 130.54, 129.90, 128.80, 128.50, 128.37, 128.14, 127.12, 126.86, 63.11, 59.36, 53.72, 53.40, 46.41, 45.83, 45.07, 39.80, 38.72, 37.38, 37.27, 35.68, 29.36, 28.92, 26.72, 22.33, 13.00, 11.53, 10.21. HRMS (ESI-TOF) Calcd for $C_{39}H_{44}ClN_8O_3S^+$ $[M+H]^+$ : 739.2940; found: 739.2956.

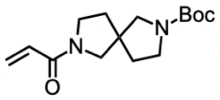

Boc
N
N
O


**5**

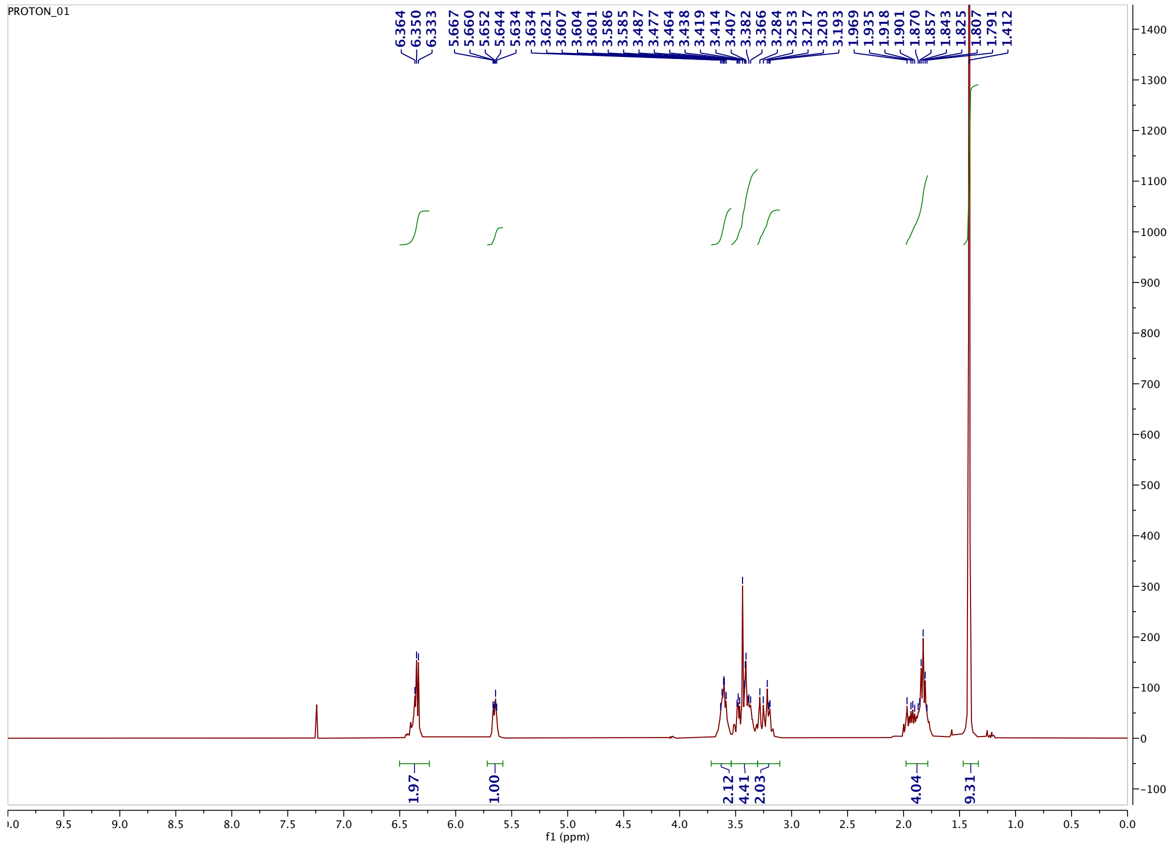

PROTON_01
6.364
6.350
6.333
5.667
5.660
5.652
5.644
5.634
3.634
3.621
3.607
3.604
3.601
3.586
3.585
3.487
3.477
3.464
3.438
3.419
3.414
3.407
3.382
3.366
3.284
3.253
3.217
3.203
3.193
1.969
1.935
1.918
1.901
1.870
1.857
1.843
1.825
1.807
1.791
1.412
1.97
1.00
2.12
4.41
2.03
4.04
9.31
f1 (ppm)


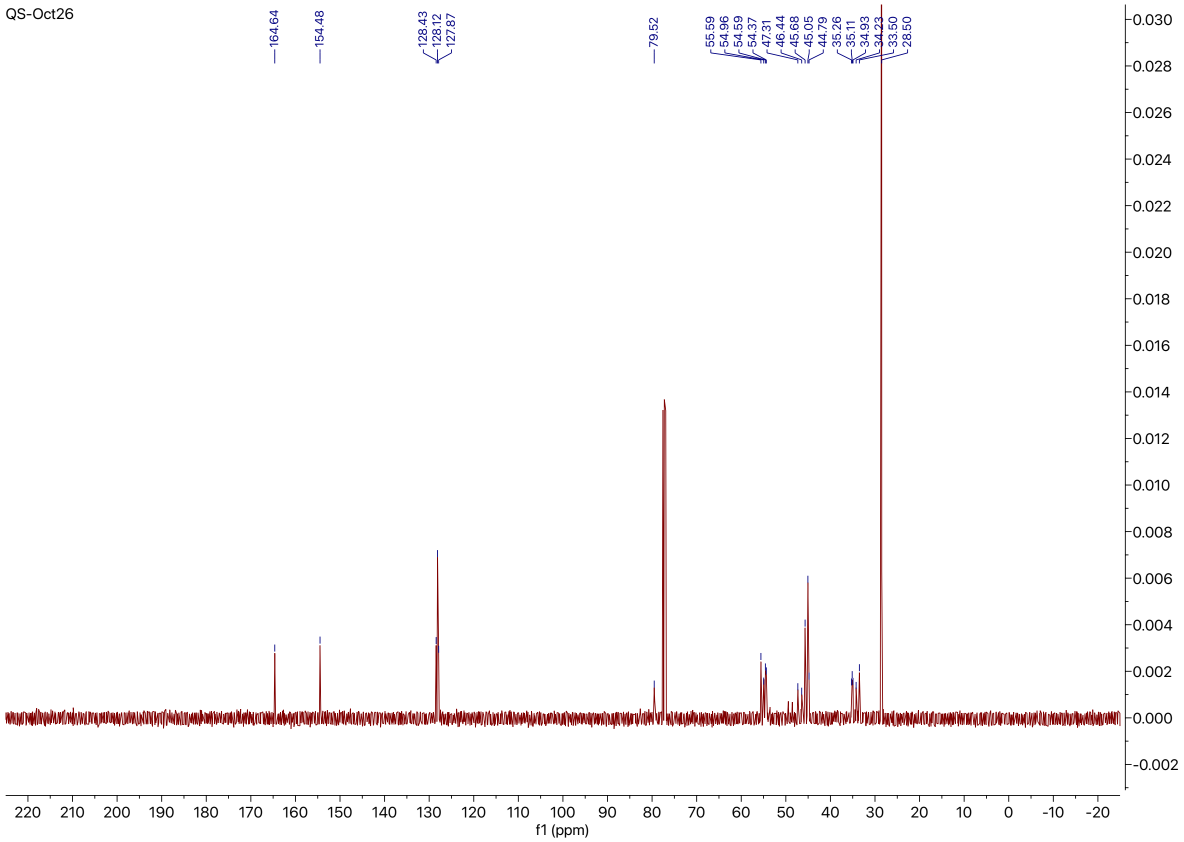

QS-Oct26
164.64
154.48
128.43
128.12
127.87
79.52
55.59
54.96
54.59
54.37
47.31
46.44
45.68
45.05
44.79
35.26
35.11
34.93
34.23
33.50
28.50
f1 (ppm)

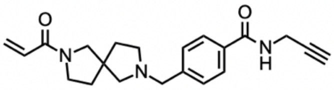

**EN-171 Alkyne**

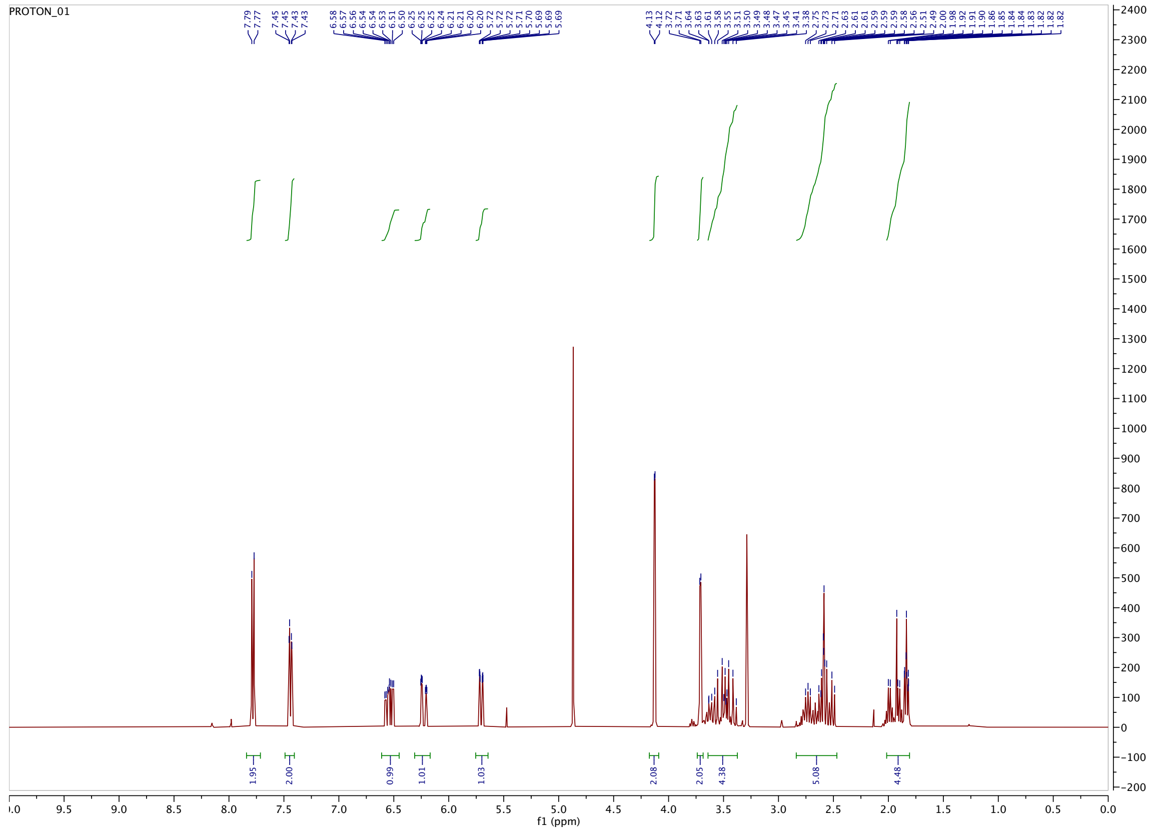


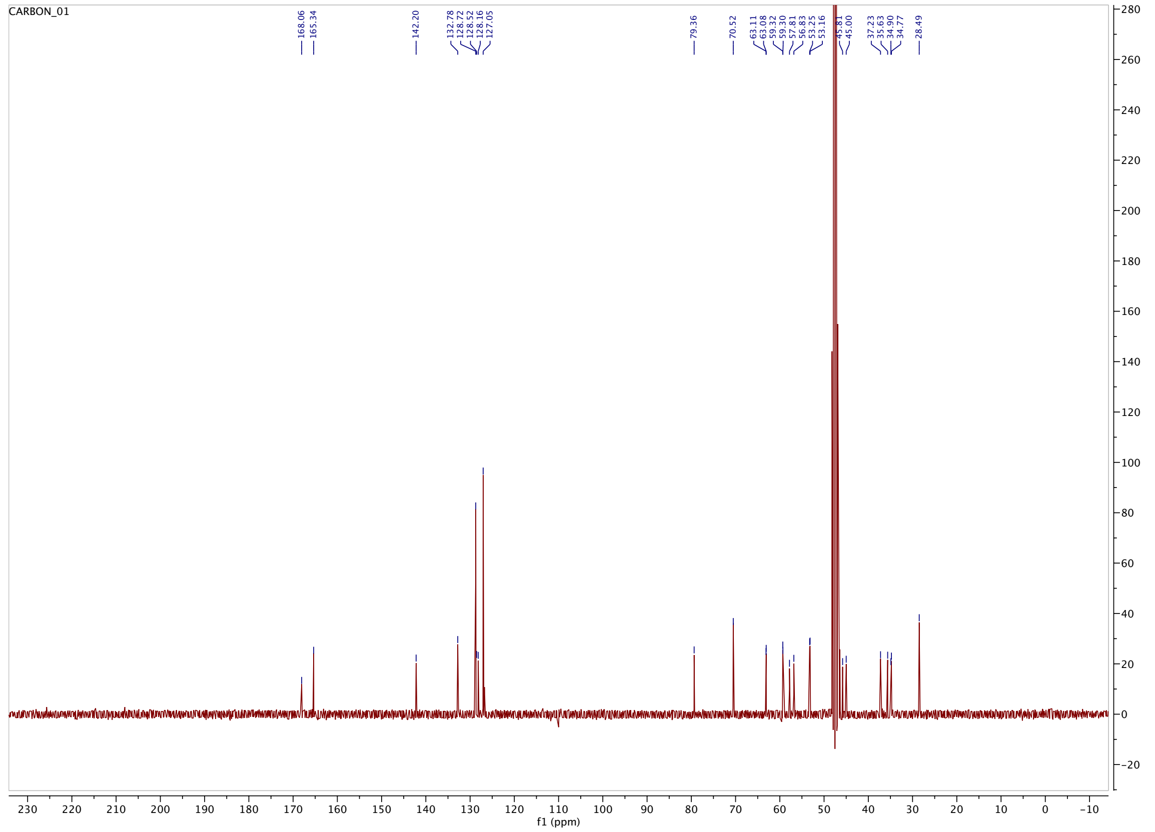

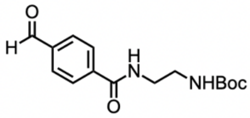
O
H
H
N
NHBoc
O

**8**

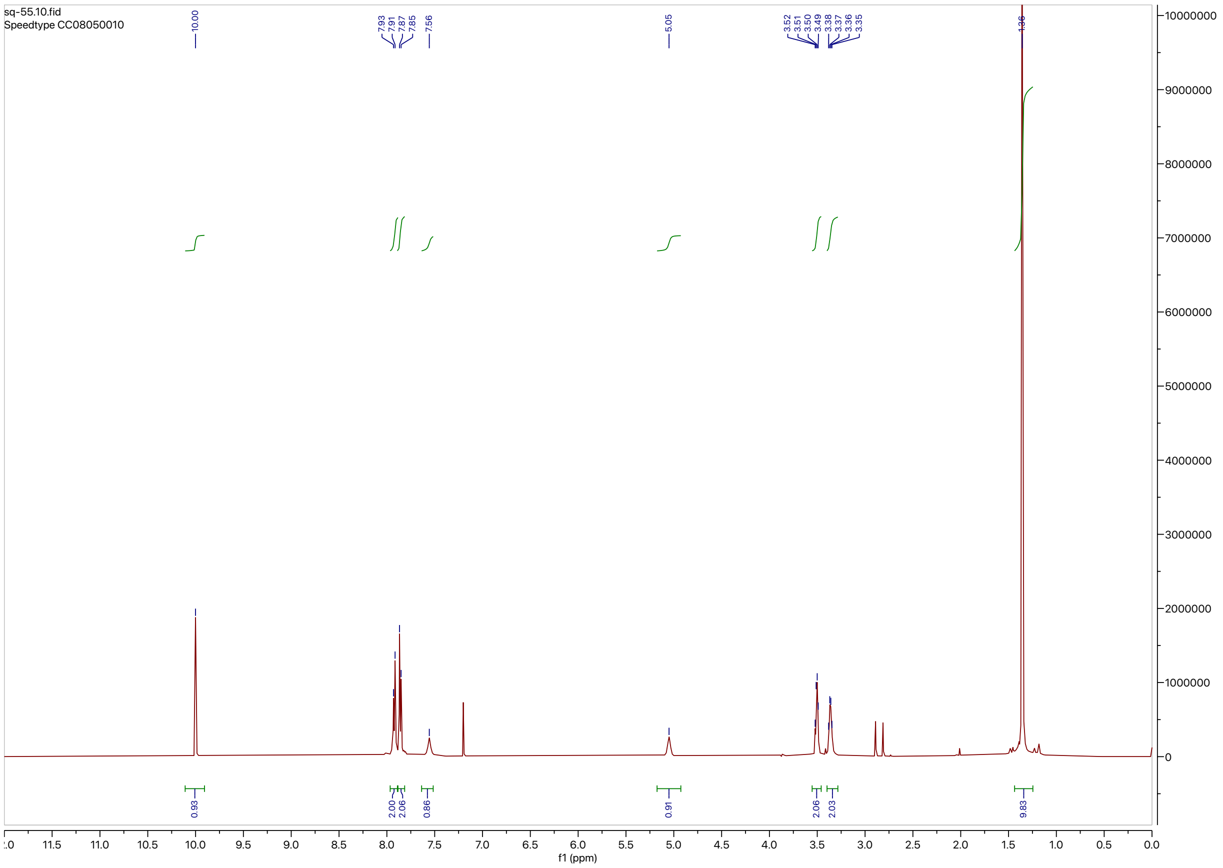
sq-55.10.fid
Speedtype CC08050010
10.00
7.93
7.91
7.87
7.85
7.56
5.05
3.52
3.51
3.50
3.49
3.38
3.37
3.36
3.35
1.36
0.93
2.00
2.06
0.86
0.91
2.06
2.03
9.83
f1 (ppm)

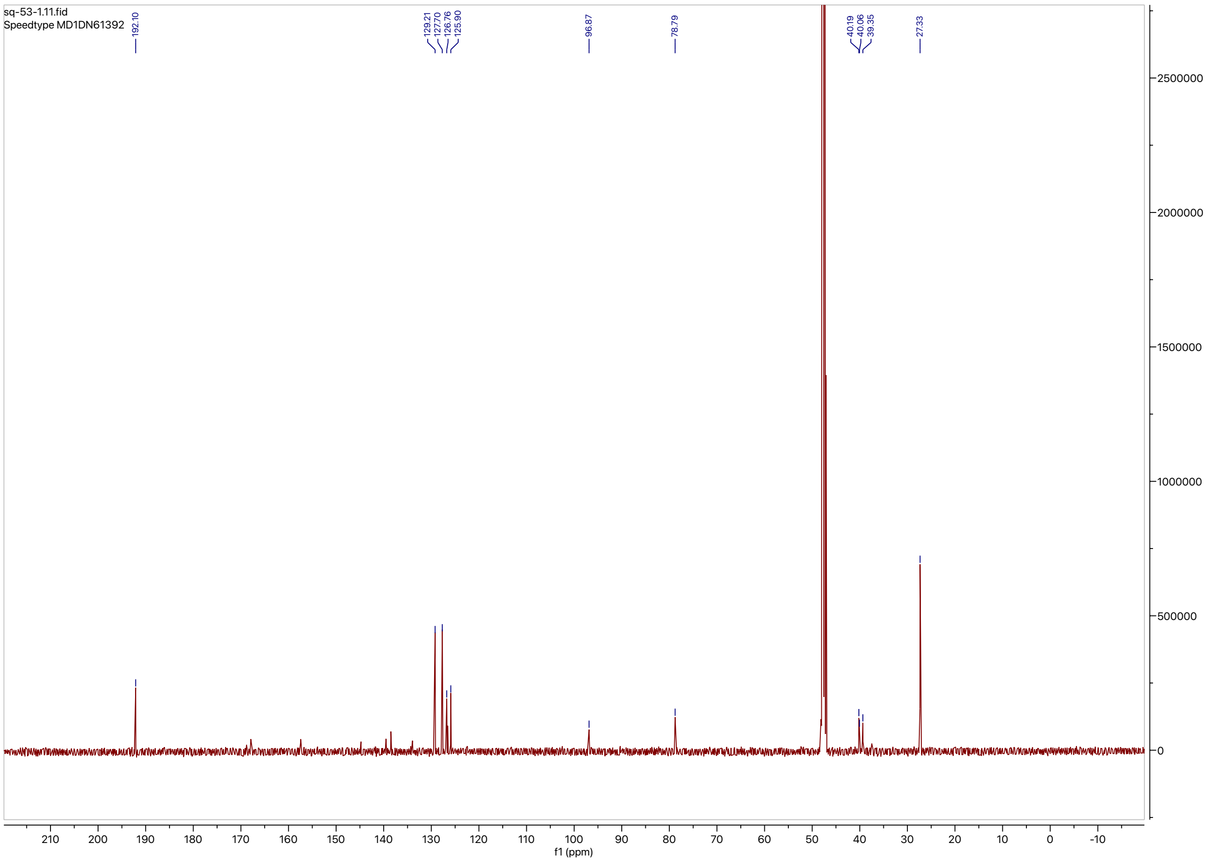
sq-53-1.11.fid
Speedtype MD1DN61392
192.10
129.21
127.70
126.76
125.90
96.87
78.79
40.19
40.06
39.35
27.33
f1 (ppm)

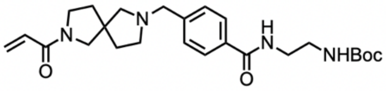

NHBoc


**9**

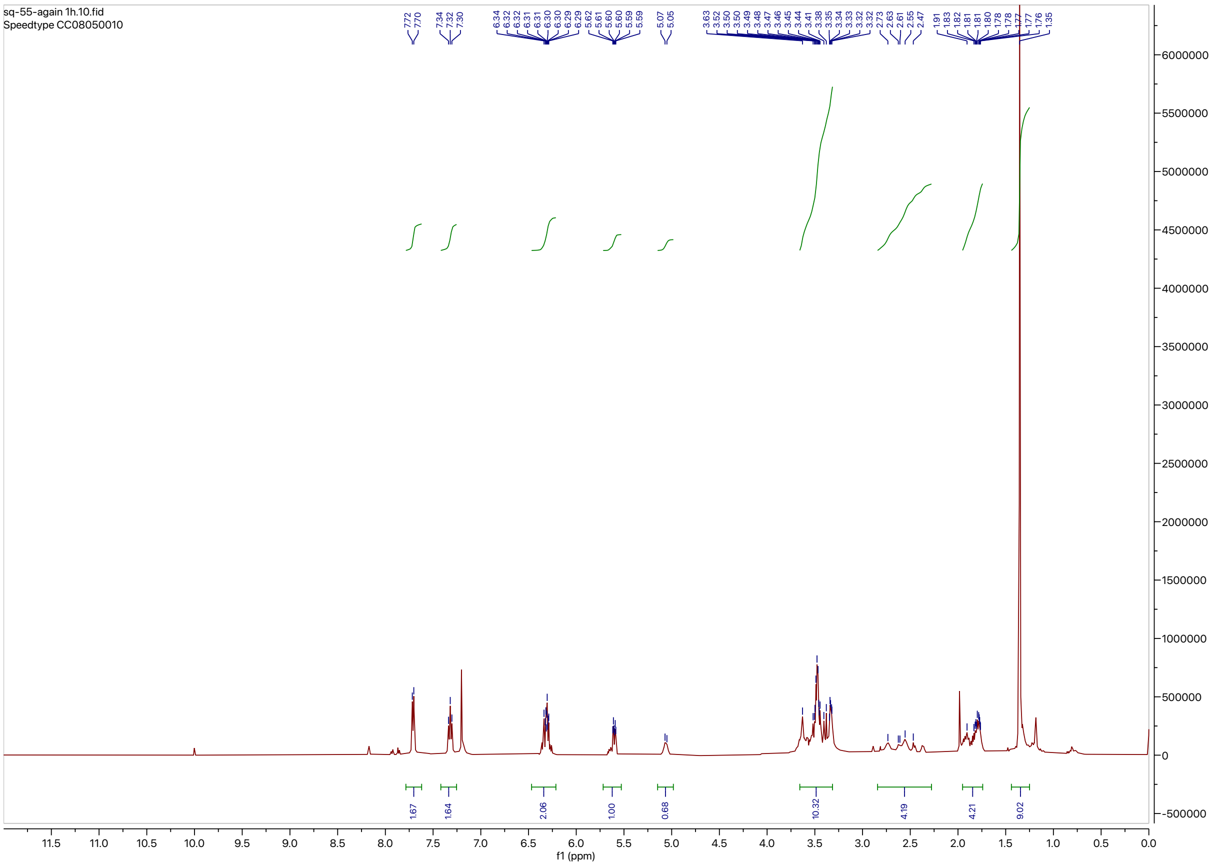

sq-55-again 1h.10.fid
Speedtype CC08050010
7.72 7.70 7.34 7.32 7.30
6.34 6.32 6.32 6.31 6.31 6.30 6.30 6.29 6.29 5.62 5.61 5.60 5.60 5.59 5.59
5.07 5.05
3.63 3.52 3.50 3.49 3.48 3.47 3.46 3.45 3.44 3.41 3.38 3.35 3.34 3.33 3.32 3.32
2.73 2.63 2.61 2.55 2.47
1.91 1.83 1.82 1.81 1.81 1.80 1.78 1.78 1.77 1.77 1.76 1.35
1.67 1.64 2.06 1.00 0.68 10.32 4.19 4.21 9.02
f1 (ppm)


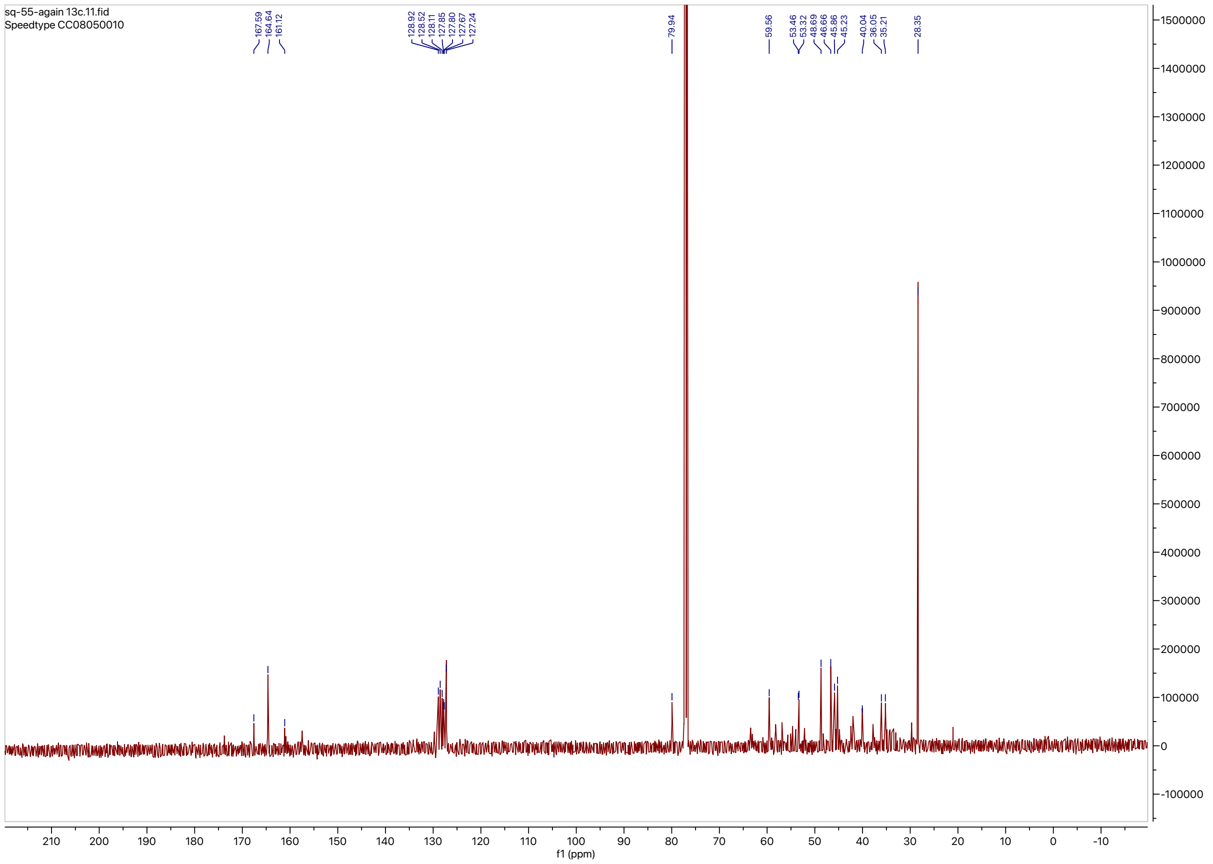

sq-55-again 13c.11.fid
Speedtype CC08050010
167.59 164.64 161.12
128.92 128.52 128.11 127.85 127.80 127.67 127.24
79.94
59.56 53.46 53.32 48.69 46.66 45.86 45.23 40.04 36.05 35.21 28.35
f1 (ppm)

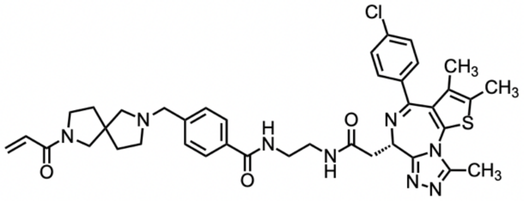
Cl
CH3
CH3
S
N
N
H
N
O
N
O
O
N
H
N
N
N
N
CH3

**QS-57**

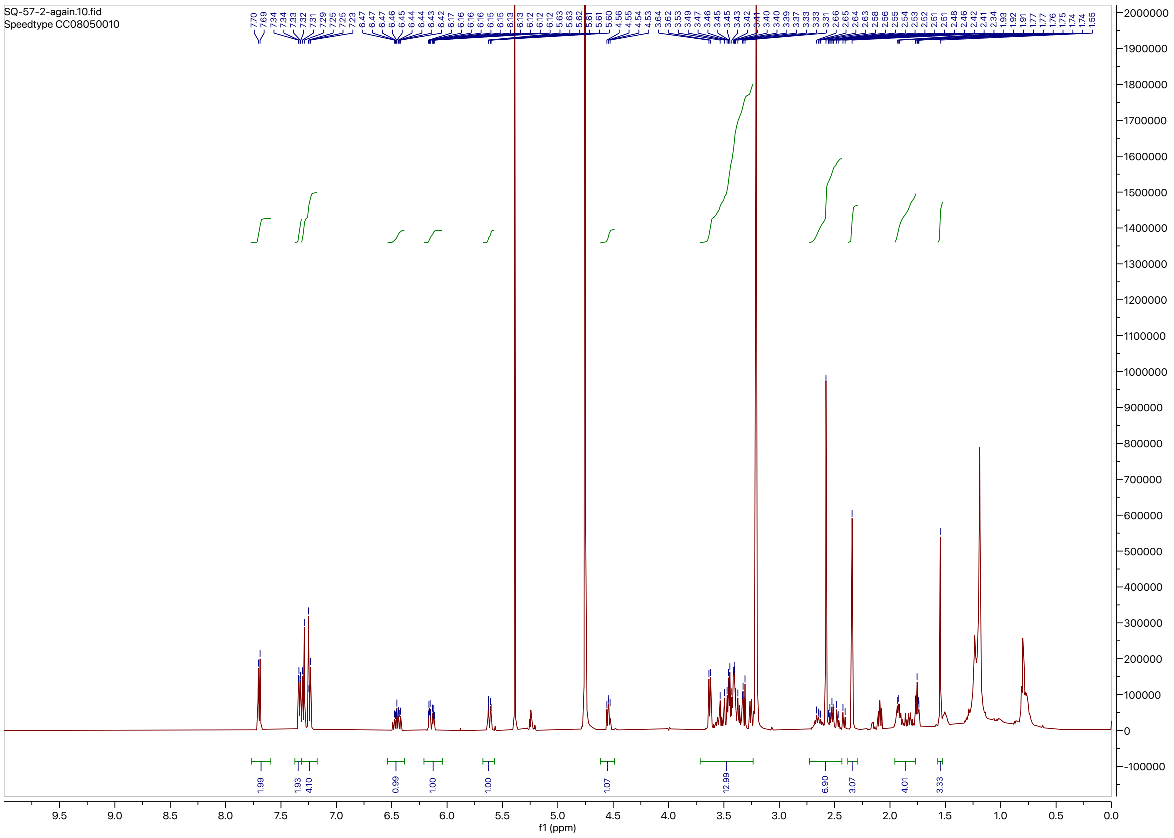
SQ-57-2-again.10.fid
Speedtype CC08050010
7.70 7.69 7.34 7.33 7.32 7.31 7.29 7.25 7.25 7.23 6.47 6.47 6.46 6.45 6.44 6.44 6.43 6.42 6.17 6.16 6.16 6.16 6.15 6.15 6.13 6.13 6.12 6.12 6.12 5.63 5.63 5.62 5.61 5.61 5.60 4.56 4.55 4.54 4.53 3.64 3.62 3.53 3.49 3.47 3.46 3.45 3.45 3.43 3.42 3.41 3.40 3.40 3.39 3.37 3.33 3.33 3.31 2.66 2.65 2.64 2.63 2.58 2.56 2.55 2.54 2.53 2.52 2.51 2.48 2.46 2.42 2.41 2.34 1.93 1.92 1.91 1.77 1.76 1.75 1.74 1.55
1.99 1.93 4.10 0.99 1.00 1.00 1.07 12.99 6.90 3.07 4.01 3.33
f1 (ppm)

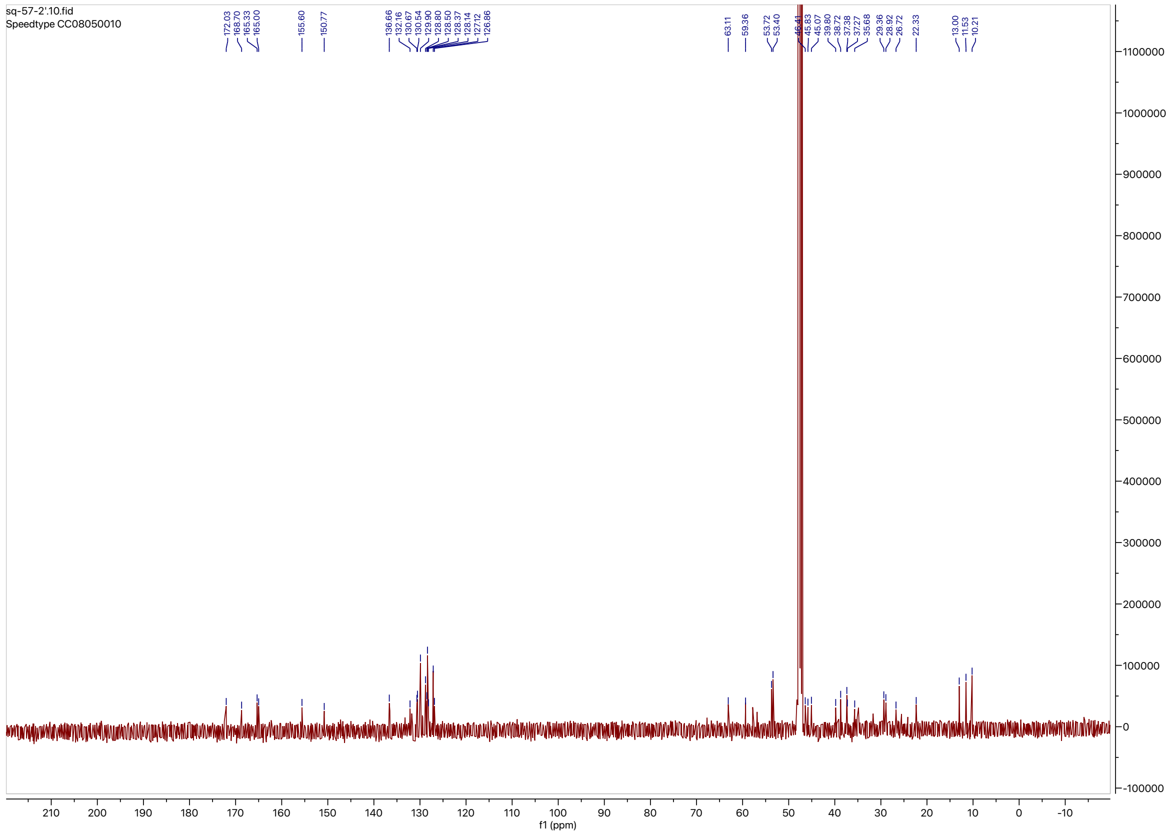
sq-57-2'.10.fid
Speedtype CC08050010
172.03 168.70 165.33 165.00 155.60 150.77 136.66 132.16 130.67 130.54 129.90 128.80 128.50 128.37 128.14 127.12 126.86 63.11 59.36 53.72 53.40 46.41 45.83 45.07 39.80 38.72 37.38 37.27 35.68 29.36 28.92 26.72 22.33 13.00 11.53 10.21
f1 (ppm)
